# Systemic and local regulation of root growth by vascular trehalose 6-phosphate is correlated with re-allocation of primary metabolites between shoots and roots

**DOI:** 10.1101/2025.11.13.688188

**Authors:** Moritz Göbel, Jacqueline Foster, Philipp Westhoff, Noemi Skorzinski, Hannah L Lepper, Maria F Njo, Markus Schmid, Tom Beeckman, Miloš Tanurdžić, Anna Amtmann, Franziska Fichtner

## Abstract

Trehalose 6-phoshpate (Tre6P) is a key signalling molecule that reflects carbon status and integrates it with developmental decision making. Tre6P has been demonstrated to regulate various developmental processes, including vegetative growth, shoot branching, flowering, and root branching. Here, we investigate how vasculature-derived Tre6P influences root system architecture by expressing heterologous Tre6P synthase (TPS) or Tre6P phosphatase (TPP) specifically in the plant vasculature. Plants with elevated vascular-derived Tre6P levels had smaller root systems, reduced sucrose levels, and lower root metabolite levels, whereas vascular TPP overexpression had the opposite effect. Using reciprocal grafting experiments, we identified shoot-derived vascular Tre6P to be the main driver of these systemic responses. In lines with increased Tre6P in the shoot vasculature, the shoot-to-root metabolite ratios were consistently increased, indicating that Tre6P modulates metabolite allocation and utilization to balance carbon partitioning between shoot and root growth. Our data further suggest that vascular Tre6P contributed to optimizing the carbon-to-nitrogen ratio to support growth and fitness. Besides this systemic function, this study shows that root-derived Tre6P also plays a critical local role in modulating root development. Collectively, our results demonstrate that Tre6P in the vasculature functions both systemically and locally to coordinate metabolic status with root growth and architecture.

## Introduction

To optimise their reproductive success – and thus their overall fitness – plants must continually adapt to a dynamic and ever-changing environment. To accomplish this, they display a remarkable amount of plasticity. This applies to their shoot and root architecture. Both are essential for plant growth with the shoot performing photosynthesis and reproduction, and the roots being essential for water and nutrient uptake and anchorage. Plants coordinate organ growth with the supply of carbon and nutrients (Kudoyarova et al., 2015, Smith and Stitt, 2007). To integrate both endogenous and exogenous signals and convert them into an appropriate developmental response, plants have developed a system of interconnected hormonal and sugar signalling pathways that respond to various stimuli (Roychoudhry and Kepinski, 2022, Sami et al., 2019, Fichtner et al., 2024, Göbel and Fichtner, 2023). Major factors modulating these dynamics are macronutrient and sugar availability of which the allocation is regulated in a systemic manner (Chen et al., 2012, Chen et al., 2016, Durand et al., 2018). Sucrose, a major product of carbon assimilation, is transported out of photosynthetic source tissues through the phloem towards actively growing sink organs. Inorganic macronutrients, however, are taken up via the roots and transported towards the shoots, where they are assimilated and used to synthesize essential metabolites like amino acids (Pratelli and Pilot, 2014, Chen et al., 2016, Chiou and Lin, 2011). Similar to sucrose, these metabolites are also distributed via the phloem (Lalonde et al., 2003, Santiago and Tegeder, 2016). Therefore, availabilities of sucrose and other metabolites like amino acids must be coordinated in the phloem and matched with sink demand.

Acting as major metabolic signal for sucrose availability, trehalose 6-phoshpate (Tre6P) is a key regulator of plant metabolism and development that coordinates metabolic status with developmental decision making (Lunn et al., 2006, Yadav et al., 2014, Figueroa and Lunn, 2016). The conceptual Tre6P-sucrose nexus model explains the homeostatic relationship between sucrose and Tre6P. According to this model Tre6P levels not only mirror sucrose levels, but changes in Tre6P levels also modulate sucrose levels thereby maintaining sucrose within an optimal concentration range for a given cell or tissue type (Yadav et al., 2014, Figueroa and Lunn, 2016, Annunziata et al., 2025).

Besides being a signal for sucrose availability, Tre6P is also the intermediate of the trehalose biosynthesis pathway. Its levels are tightly controlled by trehalose-6-phosphate synthases (TPSs), which synthesise Tre6P from UDP-glucose and glucose 6-phosphate, and trehalose-6-phosphate phosphatases (TPPs), which dephosphorylate Tre6P to trehalose (Cabib and Leloir, 1958). While there are three TPS enzymes which are expressed and catalytically active in *Arabidopsis thaliana* (TPS1, TPS2 and TPS4) (Ramon et al., 2009, Lunn, 2007), TPS1 is the only catalytically active TPS expressed throughout plant development (Fichtner et al., 2020, Fichtner and Lunn, 2021). As such, it is likely the sole producer of Tre6P after the seed stage. TPS1 is predominantly localized in the vasculature throughout the plant including shoots and roots, and in meristematic shoot tissues (Fichtner et al., 2020, Wendrich et al., 2020, Fichtner, 2025). As the vasculature is the main link between source and sink tissues, this places Tre6P synthesis and likely signalling at a highly strategic site for systemic signalling and allocation of sucrose and potentially other nutrients. For example, increasing Tre6P levels specifically in the vasculature of Arabidopsis results in transgenic plants with an early flowering and increased shoot branching phenotype (Fichtner et al., 2021, Fichtner et al., 2024). Many other important studies have focussed on the role of Tre6P in shoot development (Schluepmann et al., 2003, van Dijken et al., 2004, Satoh-Nagasawa et al., 2006, Wahl et al., 2013, Claeys et al., 2019, Ponnu et al., 2020), while much less is known about its function in the root (vasculature).

Alongside *TPS1*, several different *TPPs* show phloem-specific expression (Wendrich et al., 2020, Fichtner, 2025) and *TPPA*, *TPPB*, *TPPD*, *TPPG*, *TPPH*, *TPPI*, and *TPPJ* have been reported to be expressed in or around lateral root (LR) primordia of developing LRs (Morales-Herrera et al., 2023). *TPPB* has been shown to be expressed specifically within LR primordia during LR formation and to be a key factor in inhibiting LR formation by reducing Tre6P levels within LR primordia (Morales-Herrera et al., 2023, Morales-Herrera et al., 2024). Another study highlighted the role of *TPPI* during primary root (PR) growth showing that *tppi* mutant plants had significantly shorter primary roots (Lin et al., 2022). While the length of the meristematic zone was not affected in the *tppi* plants, the length of the elongation zone was significantly reduced (Lin et al., 2020). Transgenic lines with modified versions of the TPS1 protein also showed primary root growth defects (Fichtner et al., 2020). However, while the Tre6P levels in these lines did not directly correlate with the strength of the phenotype, these results suggest that besides LR development, Tre6P also functions in regulating primary root growth (Fichtner et al., 2020). However, the root-specific roles of Tre6P derived from the vasculature, its native site of synthesis, remain to be elucidated.

In this study, we investigated the role of vasculature-specific changes in Tre6P levels by overexpressing a bacterial *TPS* or a *TPP* from *Caenorhabditis elegans* under the control of the vasculature-specific *GLYCINE-DECARBOXYLASE P-SUBUNIT A* (*GLDPA*) promoter from *Flaveria trinervia* (Fichtner et al., 2021). Root phenotyping of these lines revealed profound, opposite effects of high and low vascular Tre6P on root growth, with low Tre6P promoting and high Tre6P inhibiting root growth. Grafting experiments revealed local and systemic effects of vascular Tre6P. Significant changes in the accumulation and allocation of many primary metabolites were recorded in response to altered vascular Tre6P levels highlighting metabolic signals as potential mediators between shoot carbon status and root growth.

## Materials and methods

### Plant material

All lines are either in the *Arabidopsis thaliana* Columbia-0 (Col-0) wild-type background and were described earlier *pGLDPA:GUS*, *pGLDPA:otsA.1*, *pGLDPA:otsA.5*, and *pGLDPA:CeTPP.4* (Fichtner et al., 2021), or are in the *tps1-1* mutant background (Eastmond et al., 2002) and were described earlier *pTPS1:otsA.2*, *pTPS1:TPS1[ΔN_2-88_]*, *pTPS1:GUS-TPS1.5*, *pTPS1:TPS-GUS.3* (Fichtner et al., 2020).

### Generation of the CRISPR-Cas9 line

The CRISPR/Cas9 mutagenesis line (*CRISPR TPS1[ΔN_5-90_]*, see figure S7 for more details), was generated as follows: A modified GreenGate system was used to assemble the vectors for CRISPR mutagenesis (Capovilla et al., 2017). Briefly, the *enhEC1.2:pEC1.1* promoter (Wang et al., 2015) was cloned into the GreenGate module A and used to drive expression of a Cas9 sequence codon-optimized for Arabidopsis (Fauser et al., 2014) in module B, followed by the *rbcs* terminator in module C (Capovilla et al., 2017).Two single guide RNAs (sgRNAs, Table S1) targeting *TPS1* were cloned into modified GreenGate vectors D and E containing U6 promoters for sgRNA expression, as previously described (Capovilla et al., 2017). The final CRISPR–Cas9 construct was assembled from these modules using standard GreenGate cloning into the pGGZ003 vector (Lampropoulos et al., 2013), with mCherry expressed in the seed (*pAt2S3::mCherry:tMAS*; Gao et al. (2016)) in module F serving as a selection marker. The N-terminal domain of the newly generated *CRISPR TPS1[ΔN_5-90_]* line was genotyped alongside the *pTPS1:TPS1[ΔN_2-88_]*, and the wild-type *TPS1* sequence using primers specified in Table S2.

### Plant growth conditions

Seeds were surface sterilised by washing them with 80% ethanol supplemented with 0.05% Tween-20 for 5 minutes, followed by 5 minutes in 100% ethanol. After drying, the seeds were spotted onto ½ MS medium (Murashige and Skoog, 1962) and stratified for at least 3 days at 4°C. Unless otherwise stated, the plates were placed vertically into 3D-printed D-Root boxes (Silva-Navas et al., 2015) without combs and grown under LED lighting in a Polyklima M3Z-TDL+ growth cabinet (Polyklima, Germany) with 16-h photoperiods (with 10 minutes of ramping light before dawn and dusk), 100 µmol m^-2^s^-1^ light intensities, 60% humidity, and 22°C day and 18°C night temperatures. The growth under altered light regimes was tested under the same conditions with 12-h, 16-h, and 24-h photoperiods. For the sucrose feeding experiment, the medium was supplemented with 1% (w/v) of sucrose. For rosette phenotyping, seedlings were grown horizontally for 2 weeks before being transferred to soil and grown in 16-h photoperiods, 150 µmol m^-2^s^-1^ light intensities, and 22°C day and 18°C night temperatures.

### Phenotyping

For root phenotyping, square Petri dishes (12x12 cm) were scanned at the time indicated in each experiment using an Epson Perfection V600 Photo flatbed scanner (Epson, Japan). Root parameters were analysed using the EZ-Rhizo2 application (https://www.psrg.org.uk/Rhizo-II.htm) (Shahzad et al., 2018). Reconstruction of averaged roots was done using the associated Root-VIS2 programme. The same scans were used to analyse the shoot diameter, while rosette phenotyping at day 21 after sowing was performed using a Sony Alpha 7 II (Sony, Japan) camera. Both datasets were analysed using ImageJ (Version 1.54p; https://imagej.net/) to determine shoot diameters and areas. Images shown are representative of replicates within each experiment.

### Metabolite analysis

Shoots and roots were harvested from ungrafted and grafted seedlings at 10 and 19 days after germination (DAG), respectively, and flash-frozen in liquid nitrogen. Whole organ plant tissue was ground to a fine powder at liquid nitrogen temperature using a ball mill and water-soluble metabolites were extracted as described in Lunn et al. (2006). Soluble sugars and starch were measured as described in (Stitt et al., 1989), and total amino acids were measured using the fluorescamine method (Bantan-Polak et al., 2001). Tre6P, other phosphorylated intermediates and organic acids were measured as described in Supporting Method S1.

### Grafting experiments

The main grafting experiments were performed following an adapted version of the Mix-and-match protocol (Vanderstraeten et al., 2022). Seeds were surface sterilised as described before, sown on ½ MS medium and stratified for 4 days at 4°C. The plates were then placed vertically into a growth cabinet running a 16/8 photoperiod with 70 µmol m^-2^s^-1^ irradiance and 21°C/18°C day/night temperatures for 3 days. For the fourth day they were transferred to 27°C (other settings unchanged). The grafting was performed by removing one cotyledon of each seedling and separating scions from rootstocks by cutting in the upper part of the hypocotyl. Afterwards, scions and rootstocks from different seedlings were aligned in petri dishes containing two wet Whatmann papers and two nitrocellulose membranes. Post-grafting, the petri dishes were sealed, and the seedlings were regenerated for 6 days in a growth cabinet with a 16-h photoperiod, a 70 µmol m^-2^s^-1^ light intensity and 21°C/18°C day/night temperatures. Successfully grafted seedlings without adventitious roots were transferred to fresh ½ MS medium and grown vertically in in 16-h photoperiods with 100 µmol m^-2^s^-1^ light intensities and 21°C/18°C day/night temperatures for 10 days. They were harvested 17 days after grafting at zeitgeber time 8 by separating the shoot and the roots following rapid quenching in liquid nitrogen for metabolite extractions, or scanned for root phenotyping 14 days after grafting. Grafting of the *pGLDPA:otsA.1* line was performed independently using a similar protocol without increasing the temperature to 27°C on day prior to grafting. Additionally, the grafting was not performed in a sterile environment, and the seedlings were instead transferred to ½ MS medium containing 0.05% plant preservative mixture (PPM; Plant Cell Technology, USA) 7 days after grafting.

### Microscopy

To assess the expression patterns of *TPS1* and expression driven by the *GLDPA* promoter β-glucuronidase (GUS) staining assays were performed as described in (Fichtner et al., 2020). Harvested seedlings were placed in a 0.1 M sodium-phosphate buffer (pH6.7), 5 mM K_3_[Fe(CN)_6_], 5 mM K_4_[Fe(CN)_6_]x3H_2_O and 1.2 mM 5-bromo-4-chloro-3-indolyl-b-D-glucuronic acid (cyclohexylammonium salt). Seedlings were vacuum infiltrated for 30 minutes and incubated overnight at 37°C in the dark. The tissue was destained by washing several times with 70% (v/v) ethanol. Stained seedlings were examined using a Nikon Eclipse Ti microscope (Nikon, Japan) fitted with a DS-Fi2 camera and operated with the NIS-Elements software. To analyse the expression patterns on transversal root sections, GUS assays were performed as previously described (Beeckman and Viane, 2000) and GUS-stained seedlings were subjected to fixation, dehydration, and embedding as previously described (De Smet et al., 2004). GUS-stained sections were imaged using an Olympus BX51 microscope (Olympus, Japan).

### Data analysis

Data plotting and statistical analysis were performed using RStudio version 2025.05.0+496 (www.rstudio.com) running R version 4.4.1 (https://cran.rproject.org/). For plotting, the packages ggplot2 or ComplexHeatmap were used. Metabolite concentrations for heatmaps were scaled by row (z-score) prior to clustering. Clustering was performed using Pearson correlation to compute the distance matrix (1 - r), followed by hierarchical clustering using Ward’s minimum variance algorithm (*ward.D2*). The resulting dendrogram was used directly for visualisation in the heatmap. Statistical analysis was performed by conducting one-way ANOVAs, followed by Fisher’s LSD post-hoc testing. To evaluate whether metabolite concentrations differ across all time points, a two-way ANOVA was performed separately for each compound and tissue type. Figures were compiled using Inkscape (Version 1.3.2 (091e20e, 2023-11-25, custom, https://inkscape.org/)).

## Results

### TPS1 is expressed throughout the phloem of the Arabidopsis root vasculature

We used TPS1-GUS and GUS-TPS1 fusion proteins expressed under the native *TPS1* promoter and in the *tps1-1* knockout background (Fichtner et al., 2020) to examine the precise expression pattern of the TPS1 protein within the root vasculature (Figures 1a, b, c, S1, S2). TPS1 was detected in the vasculature of primary and lateral roots (LR) and in the protophloem of the meristematic zone (Figure 1a). During LR development, TPS1 was detected in early stage LR primordia (LRP) up to stage III, with expression being frequently, but not always, observed in LRP of up to stage V (Figures S1b, S2c) (as defined by Malamy and Benfey (1997)). No expression could be detected in LRP of stages VI and VII. In stage VIII LRP and emerged LRs, TPS1 protein could be observed in the vasculature (Figures S1b, S2c). Using serial cross sections from the transition zone of the root meristem towards the elongation zone, we detected the TPS1 protein specifically within phloem poles, with weaker accumulation observed in parts of the procambium and pericycle (Figures 1c, S2f).

**Figure 1.**
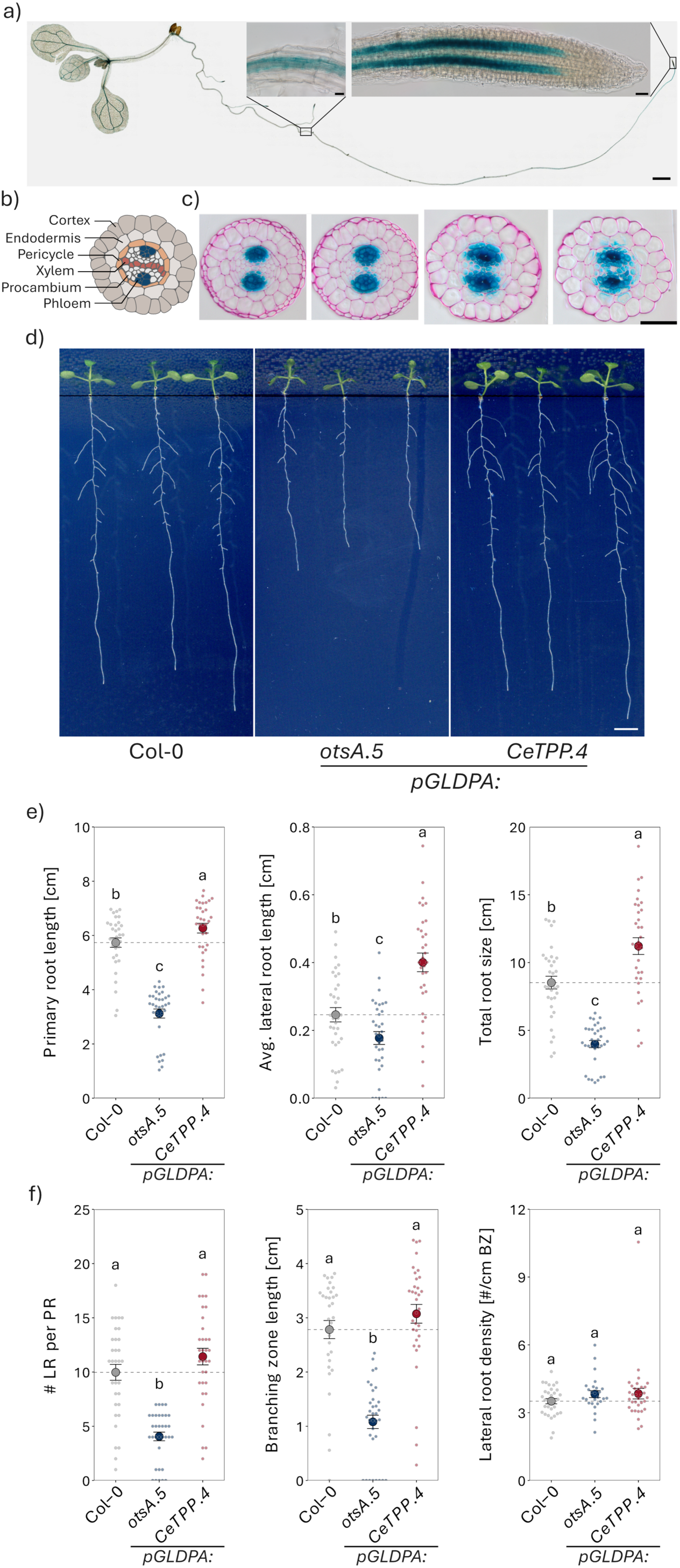
Vascular specific changes in Tre6P modify root architecture. a) A TPS1 fusion protein tagged at the C-terminus with the β-GLUCURONIDASE (GUS) reporter driven by the full length *TPS1* promoter in the *tps1-1* mutant background (*pTPS1:GUS-TPS1.5* line) was used to analyse the expression pattern of TPS1 in Arabidopsis roots 10 days after germination (DAG) (Scale bars = 1 mm for the whole seedling; 20 µM for close-up images of the root apical meristem and the vasculature). b) Schematic cross section through an Arabidopsis root. c) Transverse meristem sections of *pTPS1:GUS-TPS1.5* seedlings 10 DAG, going from the proximal meristem to the end of the root apical meristem (from left to right). Sections were counterstained with ruthenium red (Scale bar = 50 µm). c) Phenotypes of representative 10 DAG wild-type (Col-0, grey), *pGLDPA:otsA.5* (high Tre6P in the vasculature, blue), and *pGLDPA:CeTPP.4* (low Tre6P in the vasculature, red), seedlings grown vertically on ½ MS medium (Scale bar = 0.5 cm). d) Primary root (PR), average lateral root (LR), and total root sizes (sum of the PR and all LR lengths) of plants described in c). d) The number of LR per PR, the length of the branching zone (BZ) and the LR density, normalised to the BZ of plants described in c). Displayed are the mean ± SEM (large dots and error bars), with individual datapoints representing individual plants being shown as small dots (n ≥ 33). Letters indicate significant differences between genotypes according to a one-way ANOVA with *post hoc* LSD testing (*p* < 0.05).

### Vascular specific alterations in Tre6P levels resulted in root growth phenotypes

To investigate the function of Tre6P specifically within the vasculature, we used lines with alterations in Tre6P in the vasculature that were described earlier (Fichtner et al., 2021). In addition to the native Tre6P synthesis by AtTPS1 and dephosphorylation by AtTPPs, vascular tissue-specific expression of a heterologous *TPS* (*otsA* from *Escherichia coli*) or *TPP* (*CeTPP* from *Caenorhabditis elegans*) was achieved by using the *GLYCINE-DECARBOXYLASE P-SUBUNIT A* (*GLDPA*) promoter from *Flaveria trinervia* (Fichtner et al., 2021). This promoter drives vascular expression very similar to the native expression pattern of *TPS1* in the vasculature, while missing the expression in developing LRP (Figure S3). Both overexpression lines showed profound and opposite root phenotypes (Figure 1d, e, f). Compared to Col-0 wild-type plants, *pGLDPA:otsA* (line #5, always referred to if not specified otherwise) plants had shorter primary roots (PRs), reduced average LR lengths, and a reduced total root size (sum of all lengths of PR and LRs) (Figure 1e). On the contrary, *pGLDPA:CeTPP* (line #4, always referred to) plants had an increased PR length, average LR length, and total root size compared to wild-type plants (Figure 1e).

The overall decrease in PR length in *pGLDPA:otsA* plants was due to decreases in the length of both apical and branching zones (Figure S4). Similarly, the increase in PR length in *pGLDPA:CeTPP* plants coincided with an increase in the lengths of the apical and branching zones (Figure S4). *pGLDPA:otsA* plants had significantly fewer LRs per PR than wild-type plants (Figure 1f). As they also had a smaller branching zone compared to wild-type plants their LR density (as numbers of LR/branching zone length) was unchanged (Figure 1f). *pGLDPA:CeTPP* plants did not show significant differences compared to wild-type plants regarding the number or density of LRs (Figure 1f). Combined, these results indicate that vascular Tre6P influences root growth along the entire root but does not directly influence LR development, consistent with the *pGLDPA* not driving transgene expression in LRPs.

Monitoring root growth over several days confirmed these observations (Figure S5, S6). The PR and LR lengths of the *pGLDPA:otsA* plants exhibited slower growth than the wild type, reaching similar lengths as the wild type on day 10 on day 14 for PR and on day 11 for LRs (Figure S5a, S6). Further, *pGLDPA:otsA* plants showed consistently fewer LRs and a reduced branching zone length, resulting in no changes in LR density (Figure S5b). By contrast, *pGLDPA:CeTPP* plants, showed a consistent increase in PR, LR and total root length, reaching the same length about one day earlier than wild-type plants (Figure S5a). Taken together, high vascular Tre6P inhibited root growth, while lowering Tre6P in the vasculature had the opposite effect.

### Increased Tre6P levels consistently inhibit root growth

To exclude that the root growth phenotypes were caused by misexpression from the *GLDPA* promoter, we included additional lines with alterations in Tre6P metabolism. We used a second independent *pGLDPA:otsA* (line #*1*), a *pTPS1:otsA* (line #*2*), and a *pTPS1:TPS1[ΔN_2-88_]* line (Fichtner et al., 2020, 2021). The *pTPS1:otsA* and *pTPS1:TPS1[ΔN_2-88_]* lines were established previously and express either the *E. coli* TPS or a truncated version of TPS1 without the N-terminal autoinhibitory domain (missing amino acids 2 to 88, see Figure S7). The *pTPS1:otsA* line expressing a bacterial TPS results in uncontrolled synthesis of Tre6P. The N-terminal domain of TPS1 has an autoinhibitory function, deleting this domain resulted in a more active TPS1 with higher Tre6P and trehalose levels (Van Dijck et al., 2002, Fichtner et al., 2020). In addition, we generated a CRISPR-Cas9 N-terminal deletion line to avoid expression being dependent on the genomic context of the T-DNA insertion (Voichek et al., 2024, Jupe et al., 2019). In this new *CRISPR TPS1[ΔN_5-90_]* line, the amino acids 5 to 90 of the N-terminal domain were deleted, leading to the expression of a more active *TPS1* in its native locus (Figure S7). In all these lines, increased expression or activity of the various TPSs was associated with significantly decreased PR and total root lengths, supporting our previous observations using the *pGLDPA:otsA* lines, and confirming that elevated Tre6P levels within the vasculature inhibit root growth (Figure 1, 2). The strongest reduction in PR and total root length was observed in the two *pGLDPA:otsA* lines and the *pTPS1:TPS1[ΔN_2-88_]* line.

**Figure 2.**
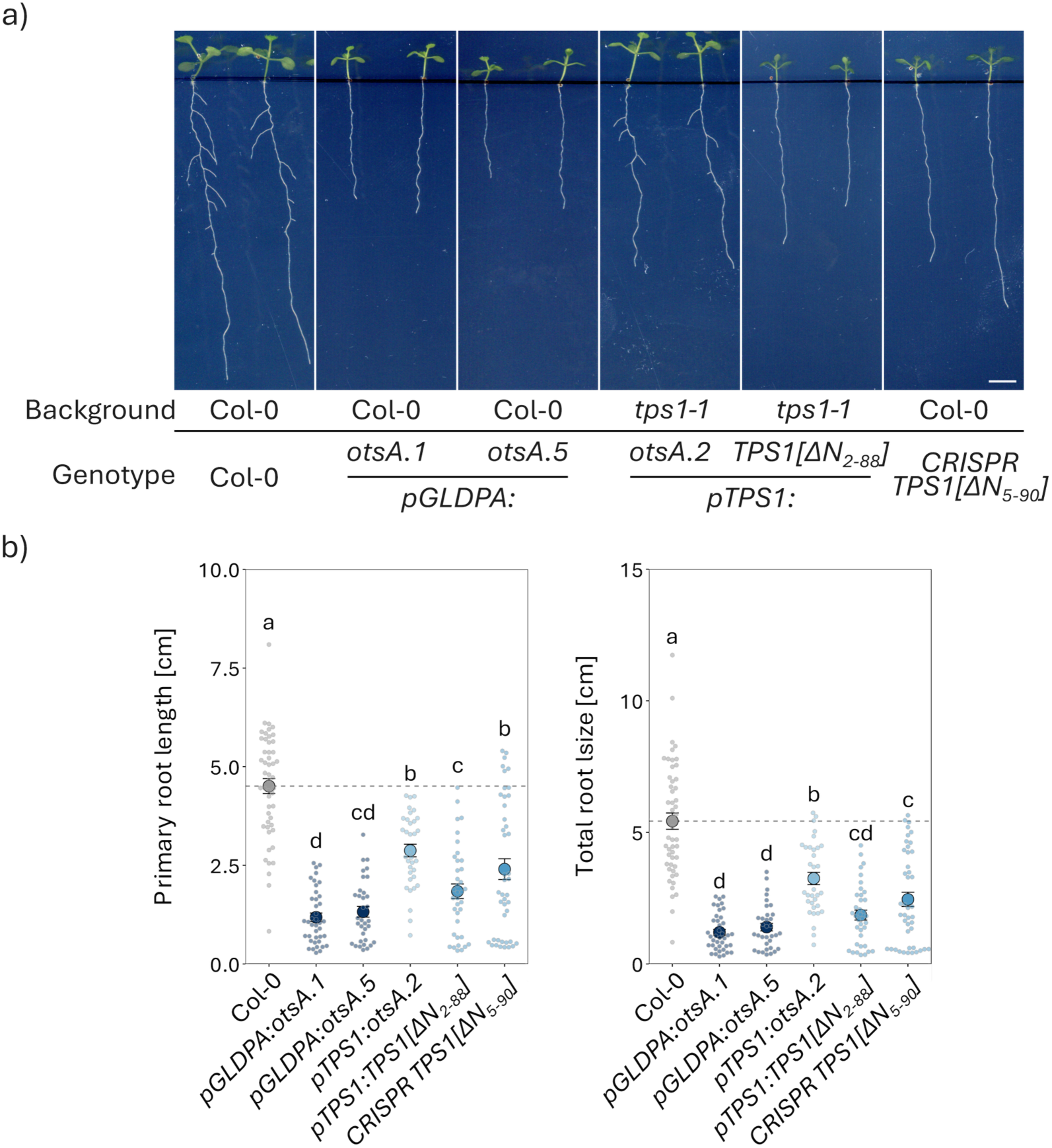
Increasing Tre6P levels by modifications in AtTPS1 also result in an inhibition of root growth. Morphological phenotypes of Arabidopsis lines expressing more active Tre6P synthase (TPS) variants. a) Phenotypes of representative 10 DAG wildtype (Col-0, grey), *pGLDPA:otsA.1* (more Tre6P in the vasculature, dark blue)*, pGLDPA:otsA.5, pTPS1:otsA.2* (expression of a heterologous TPS (otsA) under the control of the *TPS1* promoter in the *tps1-1* mutant background, light blue), *pTPS1:TPS1[ΔN_2-88_]* (expression of a TPS1 variant without the N-terminal autoinhibitory domain under the control of the *TPS1* promoter in the *tps1-1* mutant background, mid blue), and *CRISPR TPS1[ΔN_5-90_]* (removal of the TPS1 N-terminal autoinhibitory domain by CRISPR/Cas9, mid blue), seedlings grown vertically on ½ MS medium (Scale bar 0.5 cm). b) Primary (PR) and total root lengths (sum of the PR and all lateral root lengths) of the seedlings shown in a). Displayed are the mean ± SEM (large dots and error bars), with individual datapoints being shown as small dots (n ≥ 34). Letters indicate significant differences between genotypes according to a one-way ANOVA with *post hoc* LSD testing (*p* < 0.05).

### Alterations in vascular Tre6P levels modify the correlation between shoot and root growth

Given the tight coordination between root and shoot growth (Ko and Helariutta, 2017, Yang and Liu, 2020, Vercruyssen et al., 2011), we also investigated how shoot growth developed over time in the different transgenic lines, using rosette diameters as a proxy for shoot size (Figures 3a,b, S8). *pGLDPA:otsA*, *pTPS1:otsA, pTPS1:TPS1[ΔN_2-88_]*, and *CRISPR TPS1[ΔN_5-90_]* plants consistently displayed smaller rosette diameters than the wild type (Figures 3a,b, S8). Similarly, rosette diameter and area of older plants growing on soil were significantly reduced in *pGLDPA:otsA* compared to the wild type (Figure 3c,d). In contrast, the rosette diameter of *pGLDPA:CeTPP* was similar to the rosette diameter of wild-type seedlings (Figures 3b, S8), while the rosette diameter of older plants grown on soil was slightly reduced compared to wild-type plants (Figure 3c,d). However, the total rosette area was the same between *pGLDPA:CeTPP* and wild-type plants due to differences in leaf morphology (Figure 3c,d). These shoot phenotypes are both milder but consistent with those previously observed in lines overexpressing *TPS* and *TPP* under the control of a *35S* promoter (Schluepmann et al., 2003). The stronger phenotypes observed in the *35S* lines were likely caused by pleiotropic effects of expressing *TPS* and *TPP* outside the native sites of Tre6P synthesis.

**Figure 3.**
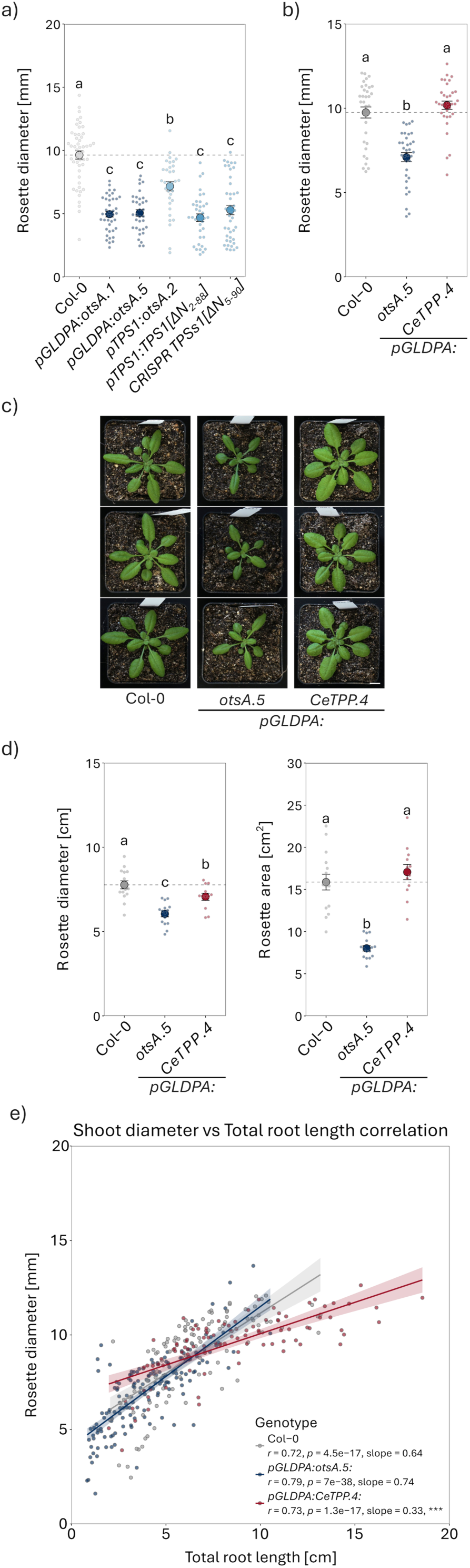
Changing vascular Tre6P levels alters rosette size and the correlation between shoot and root growth. Rosette diameter of 10 days after germination (DAG) wildtype (Col-0, grey), *pGLDPA:otsA.1* (more Tre6P in the vasculature, dark blue)*, pGLDPA:otsA.5, pTPS1:otsA.2* (expression of a heterologous TPS (*otsA*) under the control of the *TPS1* promoter in the *tps1-1* mutant background, light blue), *pTPS1:TPS1[ΔN_2-88_]* (expression of a TPS1 variant without the N-terminal autoinhibitory domain under the control of the *TPS1* promoter in the *tps1-1* mutant background, mid blue), and *CRISPR TPS1[ΔN_5-90_]* (removal of the TPS1 N-terminal autoinhibitory domain by CRISPR/Cas9, mid blue), seedlings grown vertically on ½ MS medium. b) Rosette diameter of wildtype (Col-0, grey), *pGLDPA:otsA.5* (high Tre6P in the vasculature, blue), and *pGLDPA:CeTPP.4* (low Tre6P in the vasculature, red) seedlings grown vertically on ½ MS medium (12 DAG). c) Representative images of wildtype (Col-0), *pGLDPA:otsA.5*, and *pGLDPA:CeTPP.4* plants grown in long-day conditions (16-h photoperiod) imaged 28 DAS (Scale bar = 1 cm). d) Rosette diameter and rosette area of the plants described in c). Displayed are the mean ± SEM (large dots and error bars), with individual datapoints being shown as small dots (*n* ≥ 34 for a; *n* ≥ 33 for b; n ≥ 12 for d). e) Pearson correlation analysis between the shoot diameter and total root size (sum of the PR and all LR lengths) in Arabidopsis wild-type, *pGLDPA:otsA.5*, and *pGLDPA:CeTPP.4*, seedlings grown vertically on ½ MS (all data Figures S5a and S8 are included). Genotype-specific Pearson correlation coefficients (r) and p-values (p) were calculated to assess the linear relationship between the shoot diameter and total root length. Slopes were calculated using linear regression models and are displayed with 95% confidence intervals. Significant differences between the linear relationships were assessed by fitting a linear model with an interaction between the total root size and genotype. Genotype-specific slopes were extracted using estimated marginal trends, and pairwise comparisons were performed with Col-0 as the reference (*: p < 0.05; **: p < 0.01; ***: p < 0.001).

Comparing shoot with root growth by fitting a linear model to the relationship between total root size and genotype showed that in *pGLDPA:CeTPP* plants the slope of the regression line was significantly reduced compared to the wild type (0.33 compared to 0.64; Figure 3e). This means that relative to their shoot size, the *pGLDPA:CeTPP* roots are larger, indicating that either these plants may have uncoupled their root and shoot growth responses, or that the shoot growth is more sensitive to reductions in Tre6P. For *pGLDPA:otsA* plants the correlation between shoot and root growth was not significantly different from the wild type (Figure 3e).

### Vascular Tre6P levels alter carbon allocation

The altered growth responses of the *TPS* and *TPP* overexpressing lines suggested that resources might be allocated differently when Tre6P levels in the vasculature were altered. To test this hypothesis, we determined metabolic changes in their carbon metabolism. As Tre6P is a signal for sucrose availability as well as a negative feedback regulator of sucrose levels (Lunn et al., 2006, Yadav et al., 2014), it is likely that perturbing Tre6P levels would lead to changes in sucrose metabolism. We first analysed how Tre6P and sucrose levels behaved over the course of a day in shoots and roots. Figure 4 shows the Tre6P and sucrose levels, as well as the Tre6P:sucrose ratios 8 hours after dawn (zeitgeber time, ZT, 8). In the *pGLDPA*:*otsA* line, shoot Tre6P levels were strongly increased alongside a significant reduction in sucrose content, leading to an increased Tre6P:sucrose ratio (Figure 4a). The *pGLDPA:CeTPP* line, in contrast, had higher sucrose levels, while Tre6P levels remained unchanged at ZT 8 (Figure 4a). Although not significantly different at any individual time point, Tre6P levels were significantly reduced in *pGLDPA:CeTPP* plants compared to wild-type plants based on a two-way ANOVA comparing Tre6P levels analysed across the diel cycle (*p* < 0.001). However, due to the Tre6P-sucrose nexus (Yadav et al., 2014, Annunziata et al., 2025), a decrease in Tre6P would result in accumulation of sucrose in turn resulting in activation of Tre6P synthesis, likely via activation of TPS1. Therefore, these plants likely sense and signal a continuous reduction in Tre6P, which is difficult to detect in our measurements at a single time point due to the activation of the native Tre6P synthesizing pathway. In fact, Tre6P levels in the *pGLDPA:CeTPP* line undergo minimal fluctuation over the course of the day, indicating that CeTPP is indeed constantly dephosphorylating Tre6P (Figure S9). This is further supported by the correlation between shoot sucrose and Tre6P levels. In the *pGLDPA:otsA* line, Tre6P levels vary between 0.1 and 0.3 nmol per g FW while sucrose levels did not fluctuate and were at around 0.4 µmol per g FW (Figure 4b). In *pGLDPA:CeTPP* shoots, sucrose levels fluctuated between 0.8 and 1.9 µmol per g FW, while Tre6P levels stayed relatively stable around 0.05 nmol per g FW. In both lines, the overexpression of TPS or TPP in the vasculature abolished the linear correlation between sucrose and Tre6P that was observed in wild-type shoots (Figure 4b), indicating a perturbation of the Tre6P-sucrose nexus.

**Figure 4.**
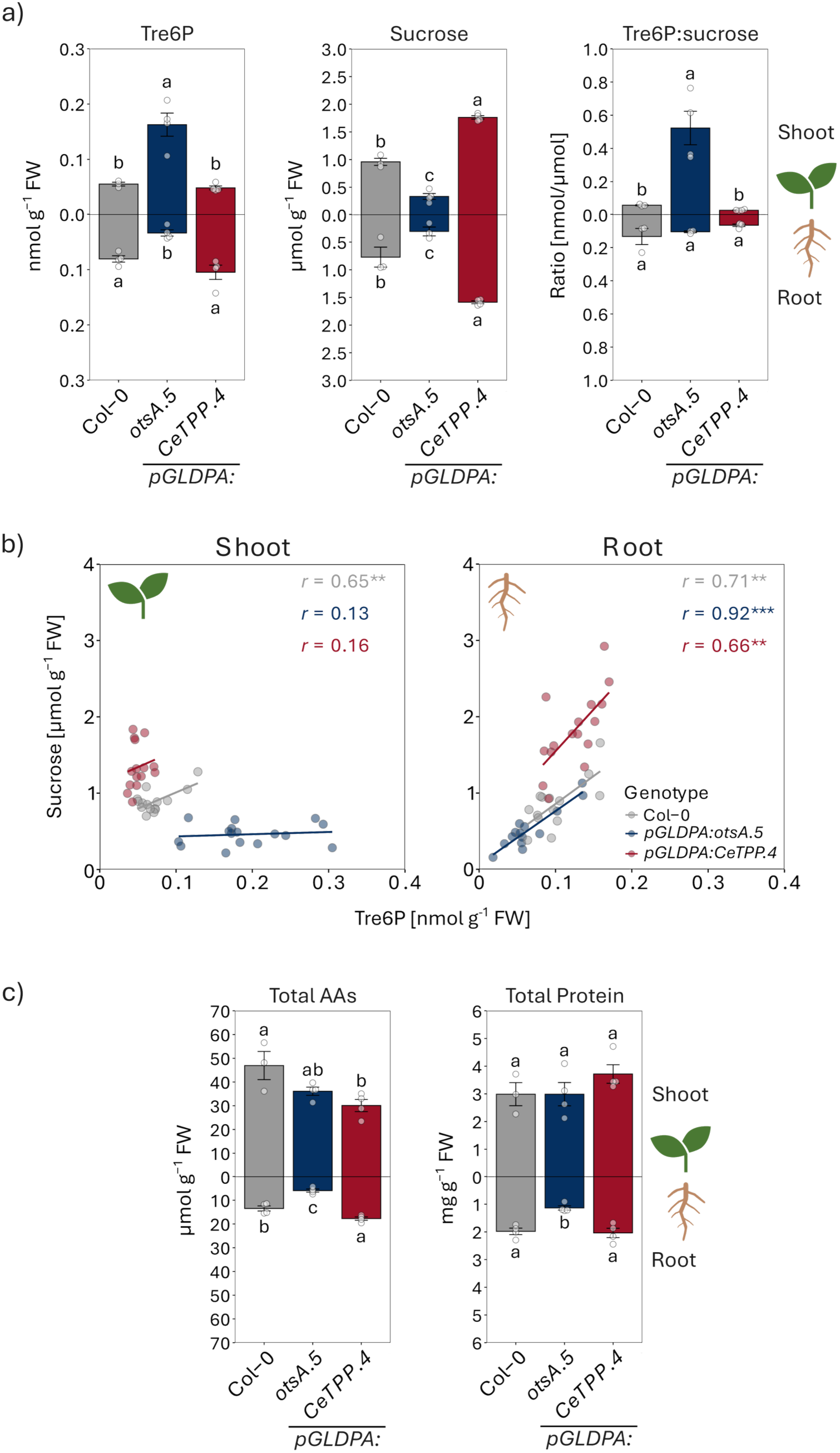
Vascular changes in Tre6P levels alter carbon allocation. Analysis of Tre6P, sucrose and other metabolites in lines expressing a heterologous TPS (otsA) or TPP (CeTPP) under the control of a vasculature-specific promoter. a) Tre6P and sucrose levels and Tre6P:ratios in shoots and roots of wild-type (Col-0, grey), *pGLDPA:otsA.5* (more vascular Tre6P, blue), *pGLDPA:CeTPP.4* (less vascular Tre6P, red) seedlings harvested 8 h after dawn (zeitgeber time (ZT) 8) from plants grown in a 16-h photoperiod (100 μmol m^−2^ s^−1^ irradiance) with 22°C day/18°C night temperatures 10 DAG. b) Pearson correlation analysis of sucrose and Tre6P data for shoots and roots harvested separately right after dawn (ZT0), 8 h after dawn (ZT8), immediately after the onset of darkness (ZT16), and right before dawn (ZT24). Genotype-specific Pearson correlation coefficients (r) and p-values were calculated to assess the linear relationship between sucrose and Tre6P levels. Significant differences between the correlations are indicated by asterisks (*: p < 0.05; **: p < 0.01; ***: p < 0.001). c) Total amino acid and total protein content of the seedlings described in a). In a) and c), values are displayed as mean ± SEM (bar graphs and error bars), with individual datapoints being shown as dots (n ≥ 3). Letters indicate significant differences between genotypes according to a one-way ANOVA with *post hoc* LSD testing (*p <* 0.05).

Tre6P levels in the roots of the *TPS* and *TPP* vascular overexpression lines did not change in the same way. The *pGLDPA:otsA* plants accumulated significantly less Tre6P based on whole root measurements, while roots of *pGLDPA:CeTPP* plants contained slightly higher Tre6P levels than wild-type plants (Figure 4a, lower panel). Sucrose levels in the roots were remarkably similar to their levels in shoots (Figure 4a, lower panel); *pGLDPA:CeTPP* plants accumulated more, while *pGLDPA:otsA* plants accumulated less sucrose compared to wild-type plants. As sucrose is supplied to the roots exclusively by photosynthetic source tissues via the phloem, root sucrose levels are expected to reflect shoot sucrose content. According to the Tre6P-sucrose nexus model, the altered root sucrose levels observed in the *pGLDPA* lines are likely to modulate Tre6P levels in the root. Consistently, reduced root sucrose levels in the roots of *pGLDPA:otsA* plants are associated with lower Tre6P levels, whereas elevated sucrose levels in *pGLDPA:CeTPP* plants coincided with higher root Tre6P levels. This is further supported by the strong positive linear correlation between sucrose and Tre6P levels in the roots of all genotypes (Figure 4b, left panel).

### Carbon supply cannot rescue the inhibition of growth in the high vascular Tre6P lines

Carbon limitation by low sucrose could potentially explain the reduced growth in lines with high vascular Tre6P (Figures 1, 2). To test this, we increased carbon supply in the *pGLDPA:otsA* line using two different approaches; sucrose feeding through the root (Liu et al., 2022, Chaudhuri et al., 2008), and increasing the photoperiod (Sulpice et al., 2014).

Wild-type plants did not show a significant change in primary root length by sucrose feeding (Figure S10a), but increasing the photoperiod significantly increased primary root length at 10 days after germination (Figure S10b). Both treatments significantly increased the primary root length of the *pGLDPA:otsA* plants (Figure S10). However, PR lengths did not reach wild-type levels irrespective of the treatment, demonstrating that increasing carbon supply and local sucrose levels alone is not sufficient to rescue the root phenotype of *pGLDPA:otsA* plants. These findings indicate that Tre6P-dependent signalling has an influence on root growth that cannot solely be attributed to alterations in sucrose allocation within the plant.

Supporting the hypothesis that sucrose allocation alone is not causing the root growth phenotypes we found that *pGLDPA:otsA* plants accumulated less total amino acids in both shoots and roots and had a reduced total protein content in roots (Figure 4c). In *pGLDPA:CeTPP*, the total amino acid content of the shoots was also significantly reduced compared to wild-type plants, but the total amino acid level of the roots was increased (Fig 4c). While roots do have the capacity to synthesise amino acids *de novo*, most amino acids are supplied to the root system by the photosynthetically active shoot, where primary nitrogen assimilation occurs and carbon skeletons are readily available from the tricarboxylic acid (TCA) cycle (Rentsch et al., 2007, Pratelli and Pilot, 2014). Like sucrose, amino acids are loaded into the phloem and transported to distant sink tissues. It is therefore conceivable that root amino acids change in a similar manner to root sucrose levels.

### Changes in Tre6P levels are accompanied by widespread changes in primary metabolism

Our data indicated that alterations in vascular Tre6P levels modified nutrient allocation from shoots to roots. To obtain more detailed information we measured a large variety of primary metabolites including soluble sugars, starch, phosphorylated intermediates, TCA cycle intermediates, and individual amino acids. Using hierarchical clustering we identified 4 clusters of metabolites behaving in a similar manner across genotypes, tissues, and ZTs (Figure 5).

**Figure 5.**
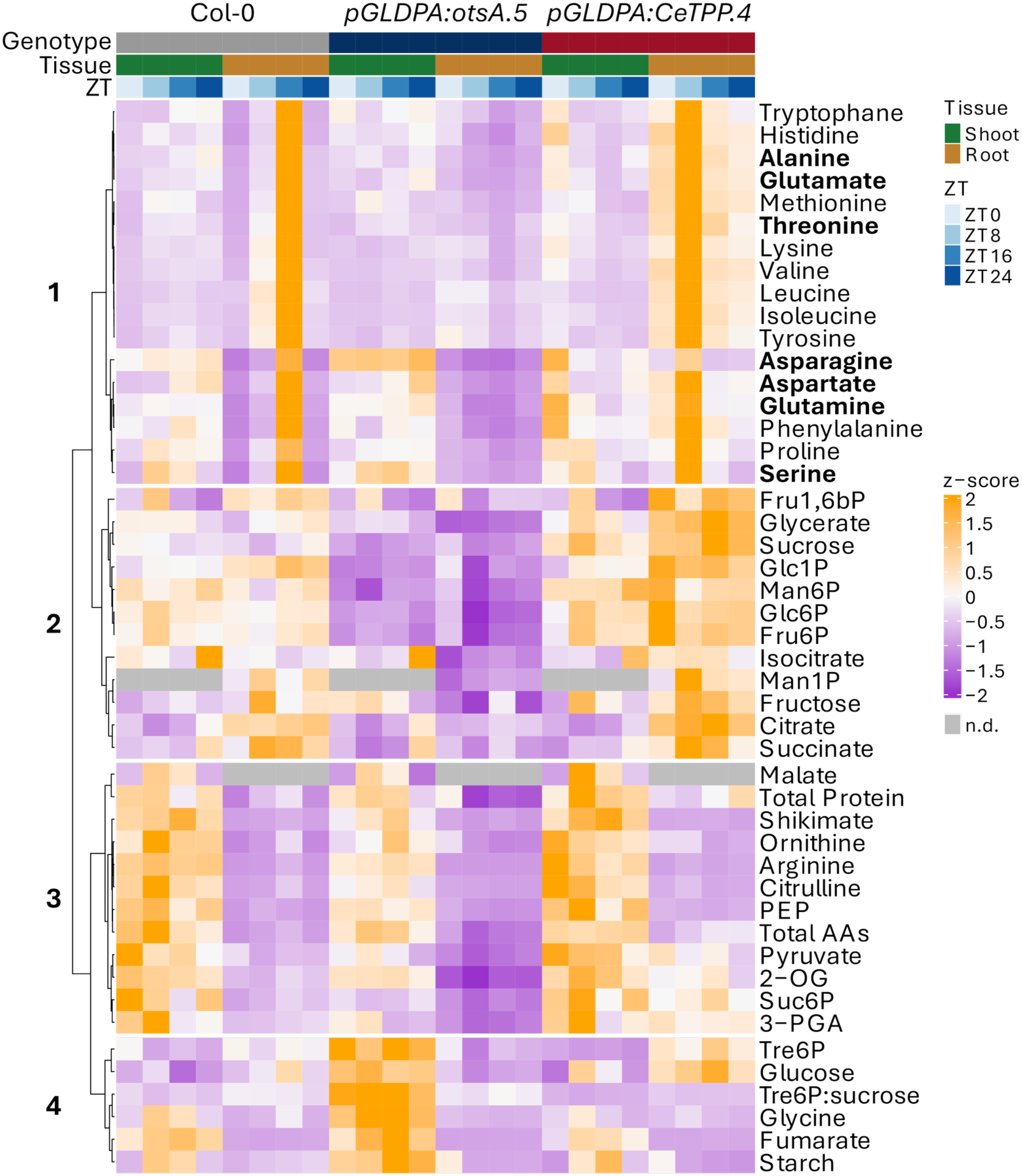
Changes in vascular Tre6P levels are accompanied by widespread changes in primary metabolism. Hierarchical clustering analysis of Tre6P and other metabolites from lines expressing a heterologous TPS (otsA) or TPP (CeTPP) under the control of a vasculature-specific promoter (data from Figures S9 and S11). Shoots (green) and roots (brown) of wild-type (Col-0, grey), *pGLDPA:otsA.5* (more Tre6P in the vasculature, blue), *pGLDPA:CeTPP.4* (less Tre6P in the vasculature, red) seedlings were harvested separately right after dawn (zeitgeber time (ZT) 0, very light blue), 8 h after dawn (ZT8, light blue), immediately after the onset of darkness (ZT16, mid blue), and right before dawn (ZT24, dark blue) from plants grown in a 16 h photoperiod (100 μmol m^−2^ s^−1^ irradiance) with 22°C day/18°C night temperatures 10 DAG. Z-scores of the mean were calculated for each metabolite and are presented as a heatmap, with high and low z-scores represented in orange and purple, respectively. Metabolites not detected in either tissue are shown in grey (n.d.). Hierarchical clustering of metabolites was performed using Pearson correlation to compute the distance matrix (1 – r), followed by Ward’s minimum variance algorithm (*ward.D2*), and is represented by a dendrogram with four distinct clusters. Amino acids most abundant in the phloem sap of Arabidopsis are highlighted.

The first cluster contains metabolites that are similar in wild-type, *pGLDPA:otsA,* and *pGLDPA:CeTPP* shoots but that are lower in *pGLDPA:otsA,* and higher in *pGLDPA:CeTPP* roots compared to wild-type roots. This cluster contains most of the proteogenic amino acids; especially the amino acids that are most abundant in Arabidopsis phloem sap, namely aspartate, glutamate, glutamine, serine, alanine, asparagine, and threonine (Besnard et al., 2016). In all three genotypes, shoot amino acid levels remained relatively constant over time (Figure S11a). In wild-type roots, amino acids accumulated over the course of the day, reaching their highest levels at dusk (ZT16) when protein synthesis is highest (Pal et al., 2013, Ishihara et al., 2015) (Figure S11b). In *pGLDPA:otsA* roots this accumulation was abolished, while *pGLDPA:CeTPP* roots contained increased levels of these amino acids and showed a shift in the accumulation pattern, reaching their peak at ZT 8 (Figures 5, S11b).

The second cluster contains metabolites that accumulated in similar amounts in shoots and roots (on a g FW basis) and that are lower in shoots and roots of *pGLDPA:otsA* plants, and higher in shoots and roots of *pGLDPA:CeTPP* plants compared to wild-type plants (Figures 5, S11). This cluster contains sucrose, phosphorylated sugars, glycerate and some TCA-cycle intermediates. Many of these compounds are primary metabolites serving as biosynthetic precursors. In both, the first and the second cluster, root metabolite levels correlate with the root phenotypes, displaying higher levels in the *pGLDPA:CeTPP* line with a larger root system, and lower levels in the *pGLDPA:otsA* line with a smaller root system.

The third cluster contains metabolites that accumulate in much higher amounts in the shoot than in the root and that are generally slightly lower in shoots and roots of *pGLDPA:otsA* plants, and marginally higher in shoots and roots of *pGLDPA:CeTPP* plants compared to wild-type plants (Figure 4, S13). This cluster included Suc6P (sucrose 6-phosphate), total amino acids, total protein content, some other amino acids, pyruvate, malate, 2-OG (2-oxoglutarate), and 3-PGA (3-phosphoglycerate).

The fourth cluster contains metabolites that accumulate more in the *pGLDPA:otsA* shoots than in *pGLDPA:CeTPP* and wild-type plants (Figure 5, S11). In this cluster, Tre6P groups together with glucose, glycine, fumarate and starch. Apart from Tre6P and glycine, all these metabolites can function as a form of carbon storage (Stitt et al., 2010, Zell et al., 2010) and are reminiscent of metabolic changes induced by increased Tre6P levels (Figueroa et al., 2016). Their accumulation in *pGLDPA:otsA* shoots is likely a direct consequence of the high Tre6P levels inhibiting starch degradation (Martins et al., 2013, Ishihara et al., 2022) and pushing carbon into other storage routes (e.g. fumarate storage in the vacuole; Figueroa et al., 2016).

Taken together, these data indicate either an altered allocation or utilization (anabolism or catabolism) of metabolites in shoots and roots induced by alterations in Tre6P levels. We therefore also explored the shoot-to-root ratios of the analysed metabolites (Figure S12). This analysis showed that while more metabolites tend to accumulate in *pGLDPA:otsA* shoots, in the *pGLDPA:CeTPP* line, metabolites preferentially accumulate in roots (Figure S12), correlating with the observed phenotypic responses (early flowering but smaller root system in *pGLDPA:otsA,* and late flowering but bigger root system in *pGLDPA:CeTPP* plants). Taken together, these findings suggest that Tre6P alters root growth by changing the allocation of metabolites between shoots and roots.

### Shoot-derived Tre6P controls root architecture systemically

To dissect the roles of shoot and root-derived Tre6P in modulating root growth we employed a grafting approach. We focused on the *pGLDPA:otsA* lines as we observed a stronger phenotype in these lines and obtained independent confirmation about the Tre6P-dependence of the phenotype in multiple lines (Figures 1, 2). As observed in ungrafted plants, *pGLDPA:otsA* (here line #*5*) homografts (*pGLDPA:otsA* scion and rootstock) exhibited the strongest inhibition of root growth, having the shortest PR and total root length among all graft combinations (Figure 6a,b). This was followed by heterografts with *pGLDPA:otsA* scions and wild-type rootstocks (Figure 6a,b), while the seedlings with *pGLDPA:otsA* roots and wild-type scions displayed the mildest inhibition in root growth (Figure 6a,b). This pattern was consistent for both PR and total root length and confirmed by a second, independent grafting experiment using the *pGLDPA:otsA* line #*1* (Figure S13). These data confirm that shoot-vasculature derived Tre6P has a stronger influence on root development than root-vasculature derived Tre6P. However, root-vasculature derived Tre6P levels still seem to have some effect on root development as expression of *otsA* in roots alone was sufficient to significantly reduce root length (Figure 6b). As observed previously, rosette diameter in grafted *pGLDPA:otsA* seedlings correlated with root length, independent of the shoot’s genotype (Figure S14). This provides further evidence that Tre6P is acting in a systemic manner to balance shoot and root growth.

**Figure 6.**
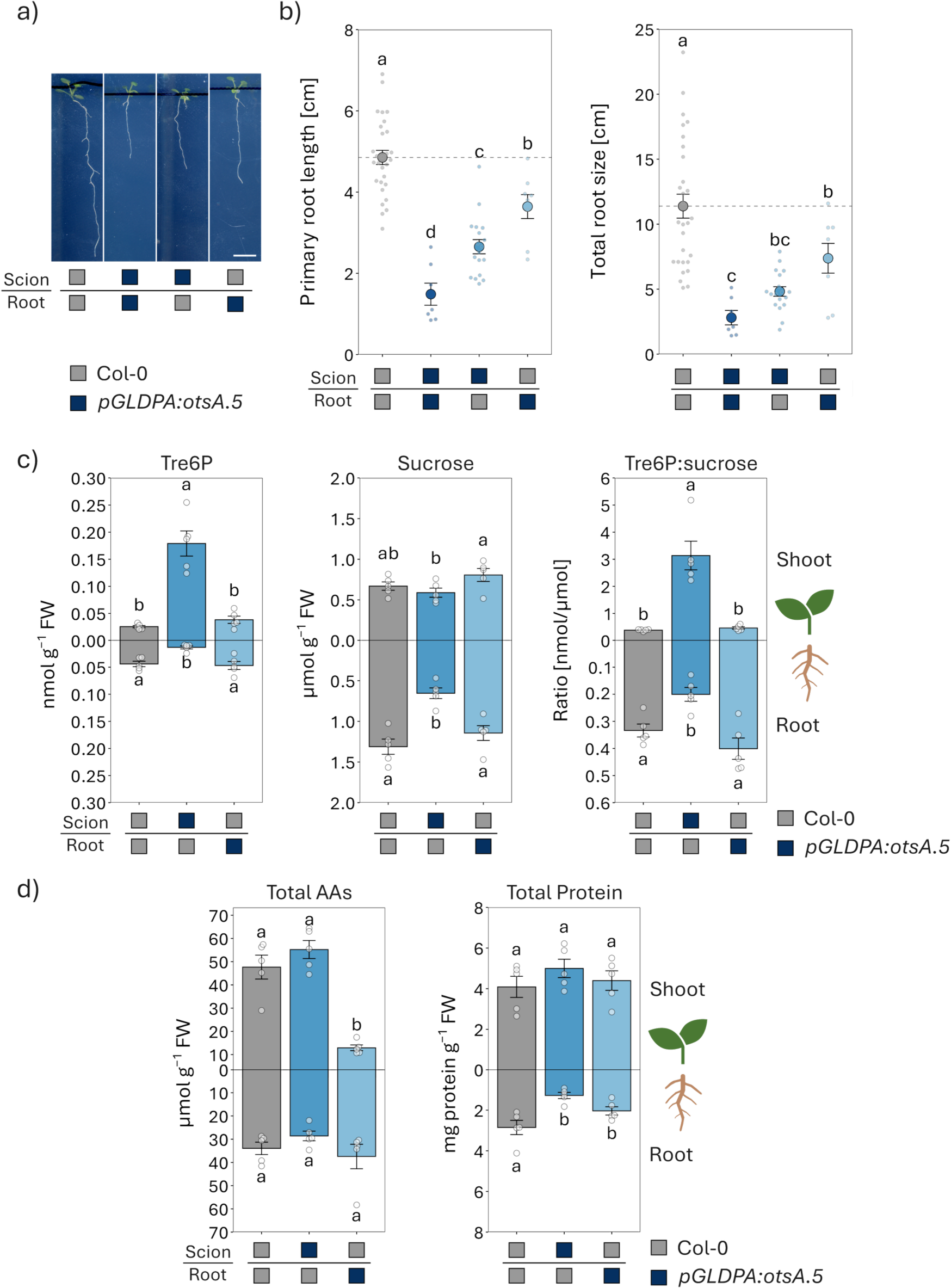
Shoot-derived Tre6P systemically controls root architecture and metabolite allocation. Root morphology and metabolite analysis of grafted Arabidopsis seedlings with root or shoot vasculature specific expression of a bacterial TPS (otsA). a) Phenotypes of representative grafted wildtype (Col-0, grey) and *pGLDPA:otsA.5* (high Tre6P in the vasculature, shades of blue represent different graft combinations) seedlings grown vertically on ½ MS medium (Scale bar 0.5 cm). b) Primary (PR) and total root sizes (sum of the PR and all lateral root lengths) of the seedlings shown in a). c) Tre6P and sucrose levels and Tre6P:sucrose ratios in shoots and roots of grafted seedlings harvested 8 h after dawn from plants grown in a 16-h photoperiod (100 μmol m^−2^ s^−1^ irradiance) with 22°C day/18°C night temperatures 19 days after germination. d) Total amino acid (AA) and total protein levels of the same seedlings. Displayed are the mean ± SEM (large dots (b)) or bar plots (c) & d)) and error bars), with individual datapoints being shown as small dots (b) n ≥ 7; c) & d) n = 5). Letters indicate significant differences between genotypes according to a one-way ANOVA with post hoc LSD testing (*p* ≤ 0.05).

Very similar to ungrafted *pGLDPA:otsA* plants, grafted seedlings with *pGLDPA:otsA* scions and wild-type rootstocks showed a strong accumulation of Tre6P in their shoots, while their root Tre6P levels were significantly reduced (Figure 6c). In contrast to ungrafted *pGLDPA:otsA* plants, sucrose levels were only reduced in the roots in the *pGLDPA:otsA* scion wild-type rootstock heterografts compared to wild-type homografts (Figure 6c). While amino acid levels were unchanged, total protein content was reduced in the *pGLDPA:otsA* scion wild-type rootstock heterografts compared to wild-type homografts (Figure 6d). In the wild-type scion *pGLDPA:otsA* rootstock heterografts, Tre6P and sucrose levels and the Tre6P:sucrose ratio were unchanged compared to wild-type homografts, however, a non-significant trend for an increase in Tre6P levels was observed in shoots (Figure 6c). These heterografts also showed a strong reduction in shoot amino acid levels and in the total protein content in the roots (Figure 6c).

When using hierarchical clustering of all measured metabolites (Figure S15, S16), three distinct clusters were identified. The first cluster contained metabolites that were increased in the *pGLDPA:otsA* scion wild-type rootstock heterografts compared to wild-type homografts. These were Tre6P, alongside a variety of metabolites spread across different pathways of the central carbon metabolism like starch, glucose, and fumarate, which all accumulated in the high Tre6P shoots (Figure S15, S16). The second cluster contained most of the sugar phosphates, which in the wild-type homografts, showed an even distribution between shoots and roots (Figure S15). This trend was abolished in seedlings with *pGLPDA:otsA* scions, where phosphorylated sugar levels were significantly reduced in the roots. The third cluster contained sucrose alongside all the measured TCA-cycle intermediates. In wild-type homografts these metabolites were more abundant in roots than in shoots. In seedlings with *pGLDPA:otsA* scions their accumulation in shoots increased, while it was decreased in roots, leading to a relatively even distribution throughout the plant (Figure S15). Therefore, accumulation of Tre6P in the shoot vasculature correlated with a consistent shift in metabolite accumulation in shoots and roots. This became more apparent when calculating the shoot to root ratios of these metabolites (Figure S17). Overall, this trend is consistent with the metabolic phenotype observed in the ungrafted *pGLDPA:otsA* seedlings, further strengthening the hypothesis that shoot-derived Tre6P is the main driver of the shift in resource allocation and the inhibition of root growth observed.

## Discussion

### Vascular Tre6P levels are an important modulator of root growth

Roots and shoots of plants have distinct roles in acquiring nutrients with roots extracting water and minerals from the soil and shoots fixing carbon from the atmosphere. Balancing these inputs is essential for optimal growth and development and requires steady communication between roots and shoots. Tre6P is a key signalling metabolite in flowering plants reporting on sucrose availability and acting as a negative feedback regulator of sucrose levels (Yadav et al., 2014, Fichtner and Lunn, 2021). This means that in response to changes in sucrose levels, Tre6P levels also change, resulting in metabolic and developmental changes, which in turn alter sucrose levels. This homeostatic relationship has been described as the Tre6P-sucrose nexus (Yadav et al., 2014, Figueroa et al., 2016). The key enzyme that synthesises Tre6P and maintains the correlation between sucrose and Tre6P in Arabidopsis plants is TPS1 (Fichtner et al., 2020). Here we showed that in Arabidopsis roots, AtTPS1 is specifically expressed in the phloem (Figure 1a,c). *AtTPPs* have a much broader expression pattern in the shoot but are predominantly expressed in the vasculature in roots (Morales-Herrera et al., 2023; Fichtner, 2025). These findings suggested an important role for Tre6P within the root vasculature.

Our biometric analysis of root phenotypes across multiple Tre6P mutants and grafts revealed consistent inverse relationship between shoot-vascular Tre6P levels and root growth. Previous studies have already suggested a role for Tre6P in root growth and development with quite different outcomes and observed phenotypes. For example, PR and lateral root growth are inhibited after a shift from normal to low light in Arabidopsis (Belda-Palazon et al., 2020). Muralidhara and colleagues, however, showed that moderate perturbations in photosynthetic activity lead to increased LR initiation and growth despite decreased sugar and Tre6P levels in the roots (Muralidhara et al., 2021). In contrast, Tre6P was found to promote lateral root growth in Arabidopsis grown in continuous light (Morales-Herrera et al., 2023). In this study, both constitutive and lateral-root-primordia-specific knockout of *AtTPPB* resulted in increased lateral root density, whereas constitutive *AtTPPB* overexpression suppressed it (Morales-Herrera et al., 2023). In contrast, *Attppi* mutants were found to have shorter primary roots and fewer lateral roots under low nitrogen (Lin et al., 2020, 2022). These data illustrate that Tre6P can have promoting and inhibiting effects, likely dependent on the location of the Tre6P pools (LR primordia, vasculature, or other tissues), the concentration range, the nutrient status, and the signalling components in specific tissues. Our results are consistent with the observed *tppi* phenotype and short-term perturbations in photosynthetic activity, suggesting that the function of Tre6P in the vasculature is different to the function of Tre6P within LR primordia.

### Root growth modulation by Tre6P might be a direct consequence of resource availability

By analysing the metabolic profile mediated by changes of Tre6P in the vasculature, we observed that carbon allocation was greatly affected (Figure 4). Besides carbon metabolism, altering Tre6P in the vasculature also had an effect on allocation of nitrogen containing metabolites like amino acids. The effect that Tre6P not only has on carbon but also on nitrogen metabolism explains why the inhibition of root growth by high Tre6P could not be rescued by increasing sugar availability (Figure S10). Vascular Tre6P levels alter the allocation and usage of carbon and nitrogen, so simply providing more carbon only partially compensates for the lack in resources.

The effect of Tre6P on altering nitrogen homeostasis might be a direct effect due to Tre6P changing transcription of amino acid transporters, or indirect by altering phloem flux and dynamics, or its influence on sucrose levels themselves. Indeed, with sucrose levels driving phloem flow (Braun et al., 2014, Ludewig and Flugge, 2013), and Tre6P influencing sucrose levels, Tre6P might indirectly control the allocation of other metabolites via altering sucrose levels and thereby mass phloem flow. This is supported by the fact that providing more carbon directly via sucrose feeding did not rescue the root phenotype of the *pGLDPA:otsA* plants (Figure S10a). The effect of a transient increase in Tre6P in Arabidopsis rosettes identified many nitrogen-associated genes to be altered (Avidan et al., 2024). The authors, however, suggested this was an indirect effect due to changes in sucrose levels, which could also contribute to the root growth phenotype in the vascular Tre6P lines analysed here. A recent study by Xia et al. supports a role for Tre6P in macronutrient signalling. By using TRAP (translating ribosome affinity purification)-sequencing to analyse gene expression in Arabidopsis companion cells under phosphate deficiency, they found *TPPJ* upregulated and *TPS5* downregulated, indicating that Tre6P levels and signalling respond to phosphate availability (Xia et al., 2025). Consistent with our results, these findings suggest that reduced Tre6P promotes root growth and that Tre6P coordinates growth with the allocation of carbon and macronutrients.

### Tre6P’s effects on root growth might be a reflection of active growth regulation through crosstalk of Tre6P with hormone signalling

Strigolactone levels are responsive to changes in nutrient availability and have been implicated in the regulation of lateral root development (Yoneyama et al., 2012, Yoneyama et al., 2015, Sun et al., 2014, Kapulnik et al., 2011, Koltai, 2011). Interestingly, we have recently shown that there is a cross talk between Tre6P and strigolactone signalling (Fichtner et al., 2024, 2025) and that strigolactone signalling is directly antagonized by sugar availability (Barbier et al., 2015, Bertheloot et al., 2020, Patil et al., 2022). Tre6P might therefore regulate root growth by directly or indirectly modifying strigolactone signalling. Potentially other hormonal signalling pathways that are influenced by sugar availability and alterations in primary metabolism (Fàbregas and Fernie, 2021), e.g. cytokinins or ABA, might contribute to the root growth modulation in the Tre6P accumulation lines.

### Conclusions

In summary, our data shows that vascular Tre6P levels modulate shoot and root growth by balancing the C/N ratio by altering the allocation and utilization of resources. Modifying Tre6P levels specifically within root and shoot tissues might therefore be a promising target for future crop breeding approaches aiming at altering shoot and root growth parameters.

## Acknowledgements

We thank Mark Stitt, John Lunn, Christine Beveridge, Guido Grossmann, and Andreas Weber for helpful project discussions, and for supporting this project. We also thank Hendrik Brodesser and Zaigham Shazad for support with plant work, and Maria Graf and Katrin Weber for technical support in the CEPLAS Metabolism & Metabolomics Laboratory. This work was supported by the German Research Foundation (Deutsche Forschungsgemeinschaft, grant number FI 2664/2-1 and FI 2664/3-1 awarded to F.F., and CEPLAS under Germany’s Excellence Strategy; project number 390686111, grant number EXC 2048/1) and the Swedish Research grant (Vetenskapsrådet, 2015-04617) awarded to M.S.

