## Supporting Information for "Systemic and local regulation of root growth by vascular trehalose 6-phosphate is correlated with re-allocation of primary metabolites between shoots and roots"

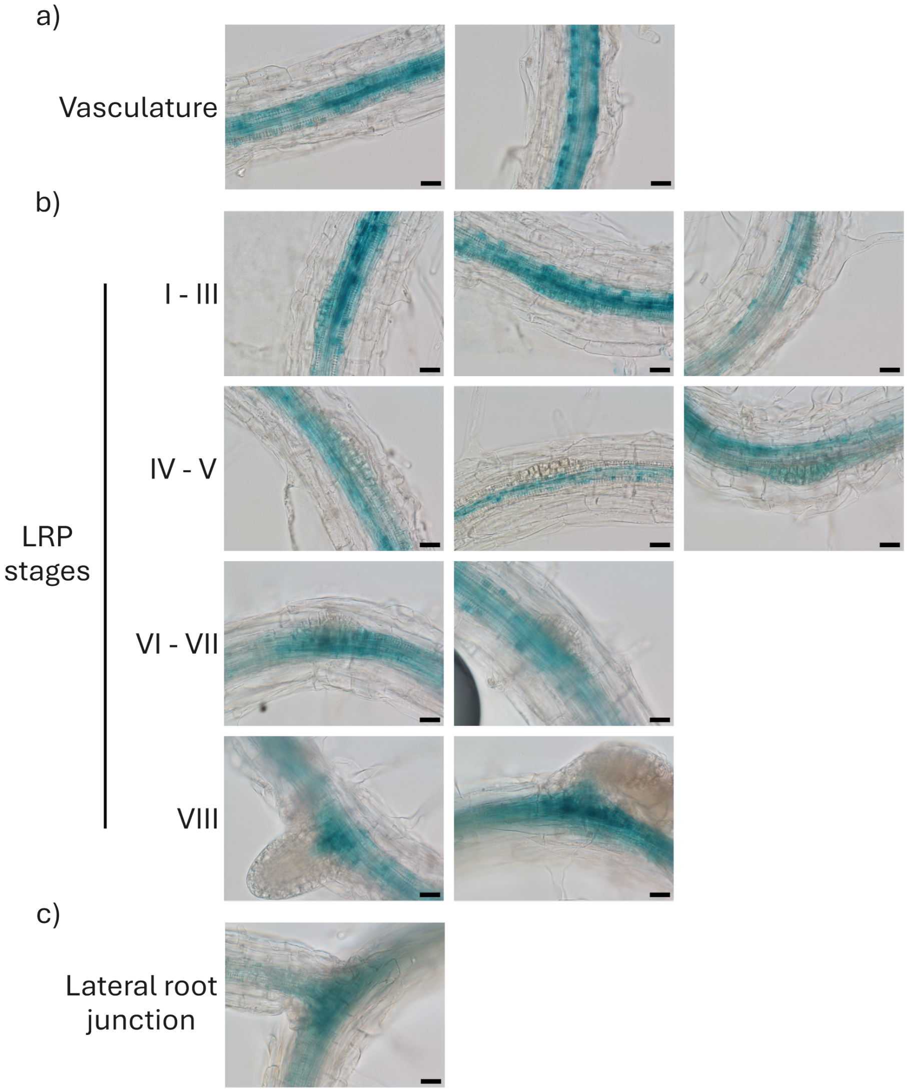


**Figure S1 Expression pattern of TPS1 in Arabidopsis roots.** A TPS1 fusion protein tagged at the C-terminus with the β-GLUCURONIDASE (GUS) reporter driven by the full length TPS1 promoter in the tps1-1 mutant background (pTPS1:GUS-TPS1.5 line) was used to analyse the expression pattern of TPS1 in Arabidopsis roots 10 days after germination. Expression a) in the vasculature, b) during lateral root emergence, c) in lateral root junctions. Scale bars = 20 µm.

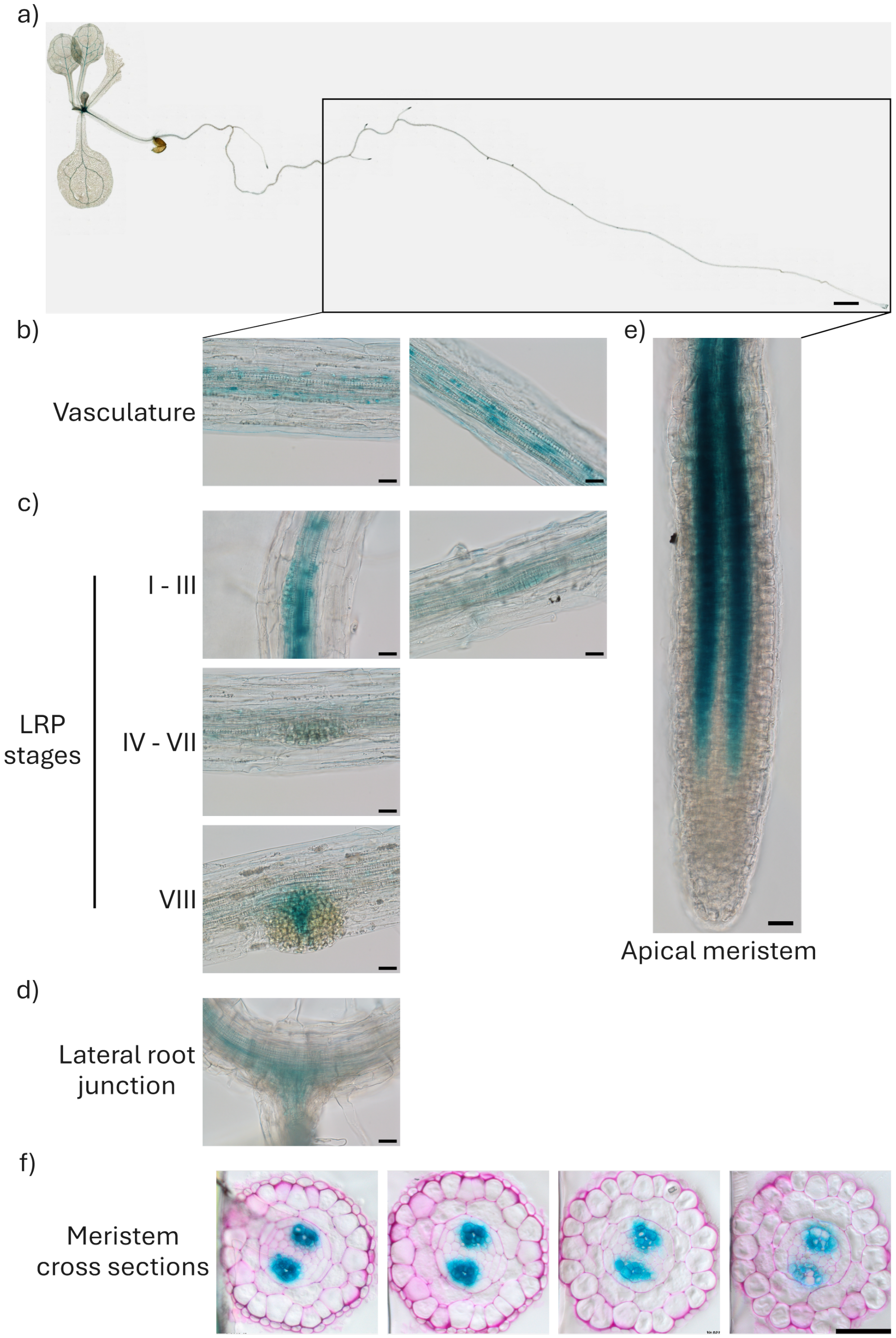


**Figure S2 Expression pattern of *TPS1* in Arabidopsis roots.** A TPS1 fusion protein tagged at the N-terminus with the β-GLUCURONIDASE (GUS) reporter driven by the full length *TPS1* promoter in the *tps1-1* mutant background (*pTPS1:TPS1.GUS.3* line) was used to analyse the expression pattern of *TPS1* in Arabidopsis roots 10 days after germination (DAG). Expression a) in the whole seedling, b) in the vasculature, c) in different stages of lateral root primordia (LRP) during lateral root emergence, d) in lateral root junctions, and e) in the apical meristem (Scale bars = 1 mm for the whole seedling; 20 µM for close-up images of the root apical meristem and the vasculature). f) Transverse sections of roots from *pTPS1:GUS-TPS1.5* seedlings 10 DAG, from the elongation zone (left) to the root apical meristem (right). Sections were counterstained with ruthenium red (Scale bar 50 µm).

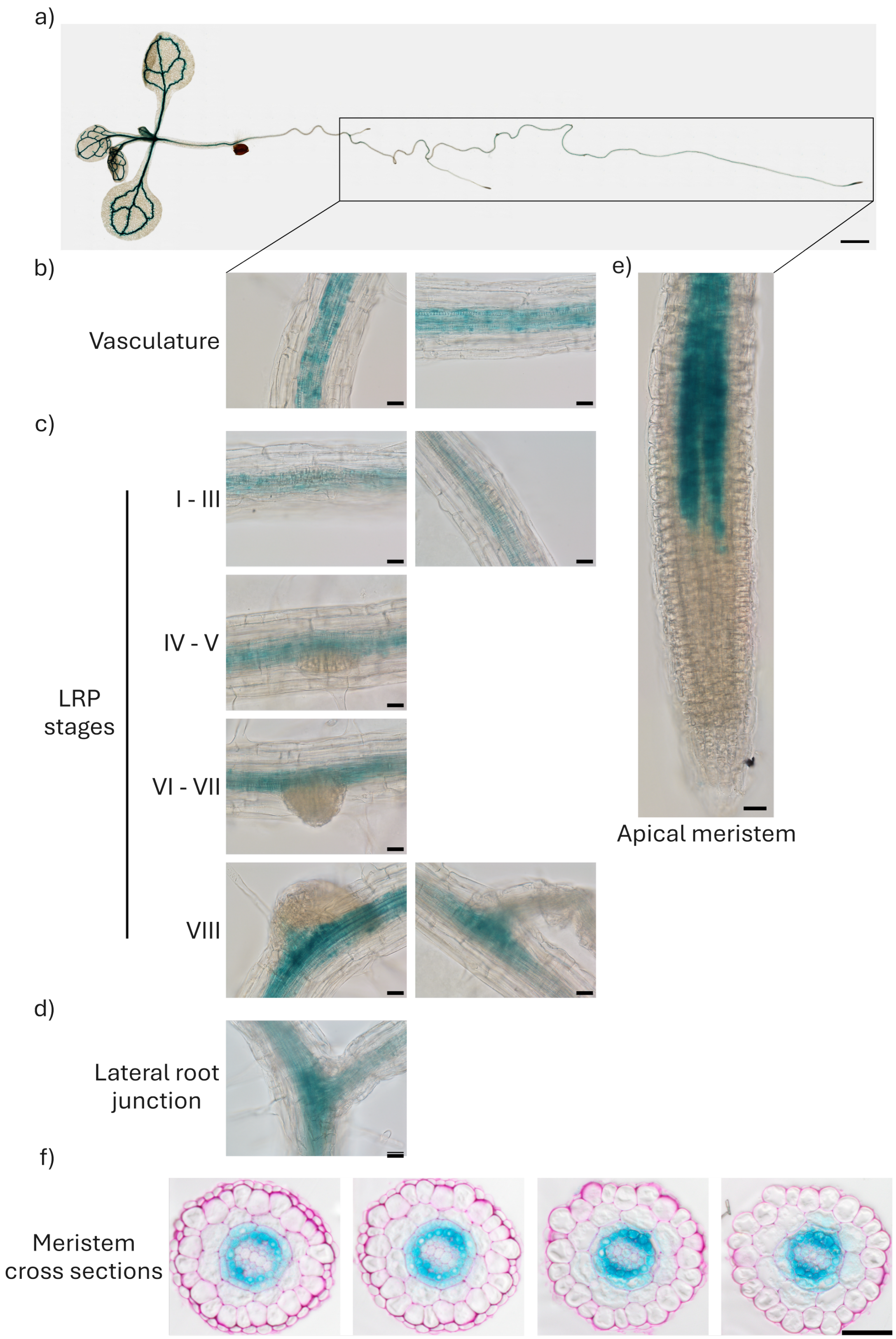


**Figure S3 Expression pattern mediated by the *GLYCINE-DECARBOXYLASE P-SUBUNIT A* (*GLDPA*) promoter from *Flaveria trinervia* in Arabidopsis**. The β-GLUCURONIDASE (GUS) reporter was expressed under the control of the *GLDPA* promoter from *Flaveria trinervia* to analyse its expression pattern in Arabidopsis roots 10 days after germination (DAG). Expression a) in the whole seedling, b) in the vasculature, c) in different stages of lateral root primordia (LRP) during lateral root emergence, d) in lateral root junctions, and e) in the apical meristem. (Scale bars = 1 mm for the whole seedling; 20 µM for close-up images of the root apical meristem and the vasculature). f) Transverse sections of roots from *pGLDPA:GUS* seedlings 10 DAG, from the elongation zone (left) to the root apical meristem (right). Sections were counterstained with ruthenium red (Scale bar 50 µm).

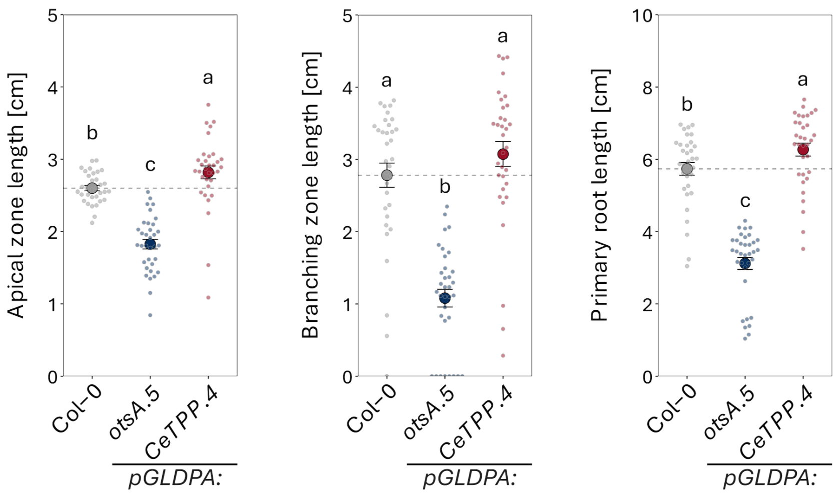


**Figure S4 Changing vascular Tre6P levels alters the root system architecture.** Root system architecture of wild-type (Col-0, grey), *pGLDPA:otsA.5* (high Tre6P in the vasculature, blue), and *pGLDPA:CeTPP.4* (low Tre6P in the vasculature, red), seedlings grown vertically on ½ MS. Supporting Figure 1 b). Depicted are the apical zone (AZ) length, the branching zone (BZ) length and the primary root (PR) length. The BZ and PR length' were duplicated from Figure 1 for ease of comparison. Displayed are the mean ± SEM (Large dots and error bars), with individual datapoints being shown as small dots ($n \geq 33$). Letters indicate significant differences between genotypes according to a one-way ANOVA with *post hoc* LSD testing ($p < 0.05$).

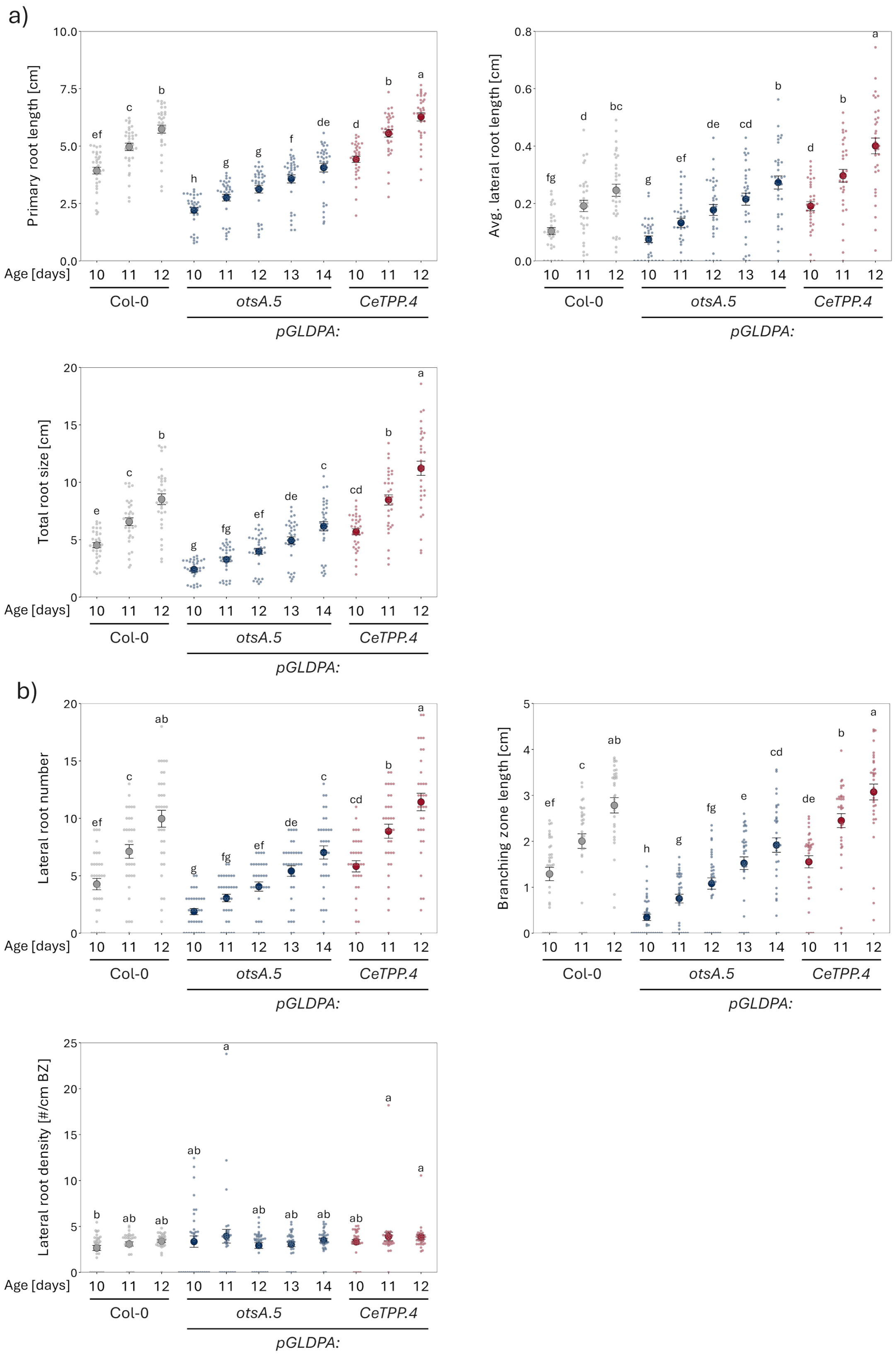


**Figure S5 Tre6P consistently alters root development.** Root system architecture of Arabidopsis wild-type (Col-0, grey), *pGLDPA:otsA.5* (high Tre6P in the vasculature, blue), and *pGLDPA:CeTPP.4* (low Tre6P in the vasculature, red), seedlings grown vertically on ½ MS medium. Primary root (PR), average lateral root (LR), total root sizes (sum of the PR and all LR lengths) (a), and LR number, branching zone length, and LR density (b) scored over time (age [days] = DAG). Displayed are the mean ± SEM (Large dots and error bars), with individual datapoints being shown as small dots ($n \geq 33$). Letters indicate significant differences between genotypes according to a one-way ANOVA with *post hoc* LSD testing ($p < 0.05$).

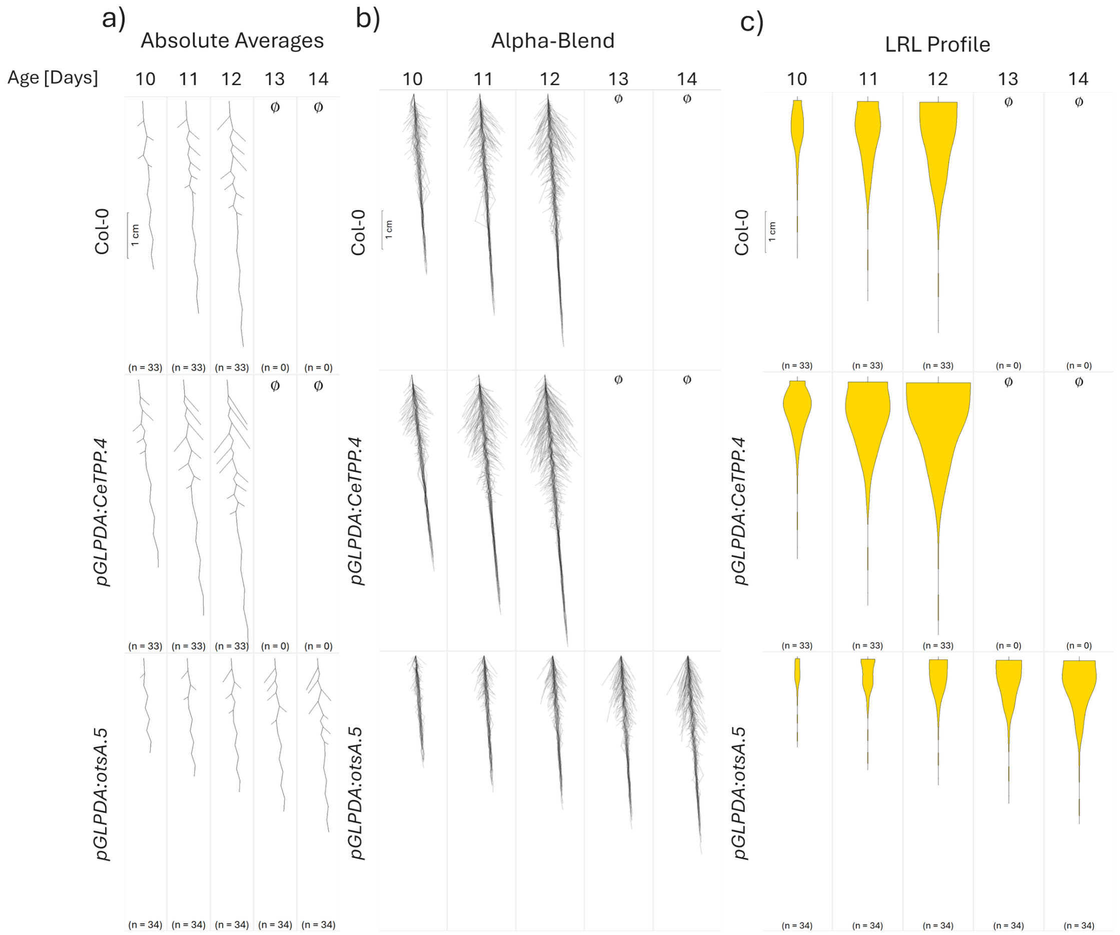


**Figure S6 Visual representation of the average root system architecture (RSA) in Arabidopsis plants with different vascular Tre6P levels.** Average root system architecture of Arabidopsis wild-type (Col-0, grey), *pGLDPA:otsA.5* (high Tre6P in the vasculature, blue), and *pGLDPA:CeTPP.4* (low Tre6P in the vasculature, red), seedlings grown vertically on ½ MS medium over time (Age [Days] = Days after germination) using the Root-VIS software. The data used for the RSA reconstructions are the same that were used in Figure 1 and S4. a) Absolute averaged RSA reconstruction. b) Alpha blended visualisation of individual replicate roots for each condition. c) Lateral root (LR) profile displayed as mean LR length (LRL) in 10 sectors across the primary root. For a) and c), the replicate number of each condition is shown below the reconstructed roots.

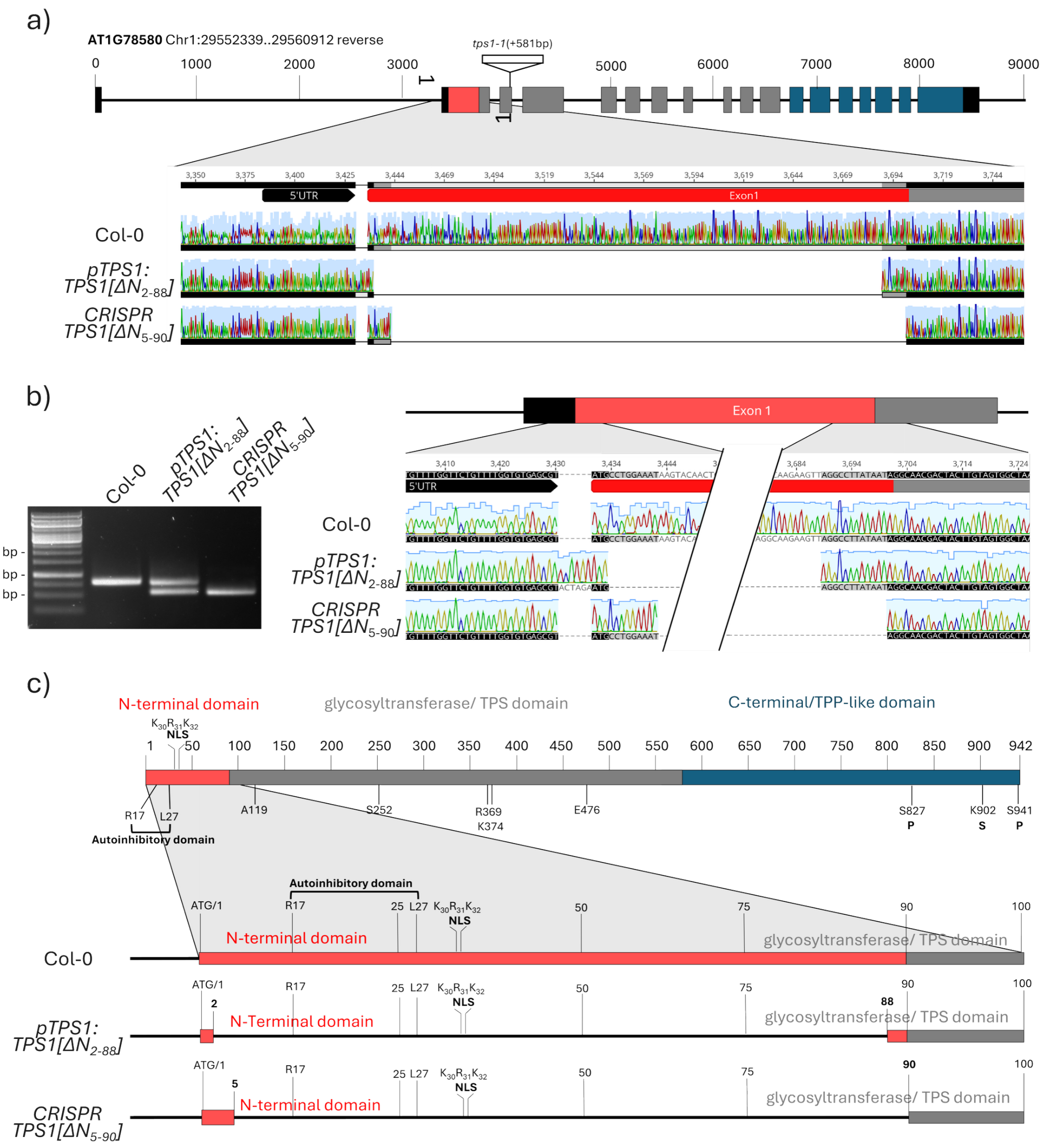


**Figure S7 Confirmation of the genotypic background of the *CRISPR TPS1[ΔN$_{5\text{-}90}$]* line.** a) Schematic overview and sequencing results showing the deletions in the N-terminal domain of TPS1 in the TPS1[ΔN$_{2\text{-}88}$] line in the *tps1-1* mutant background (Fichtner et al., 2020), and the *CRISPR* TPS1[ΔN$_{5\text{-}90}$] line compared to wild-type TPS1. b) Genomic PCR analysis confirming deletion of the N-terminal domains. n.b., the *pTPS1:TPS1[ΔN$_{2\text{-}88}$] line* was generated in the *tps1-1* knockout, in which *TPS1* is disrupted by a transposon insertion (see Fichtner et al., 2020). As the insertion lies in the TPS domain downstream of the used primers, the N-terminal domain of the native TPS1 is still present and amplified alongside the reintroduced *TPS1[ΔN$_{5\text{-}90}$]* . c) Schematic overview of the protein structures derived from the N-terminal deletions.

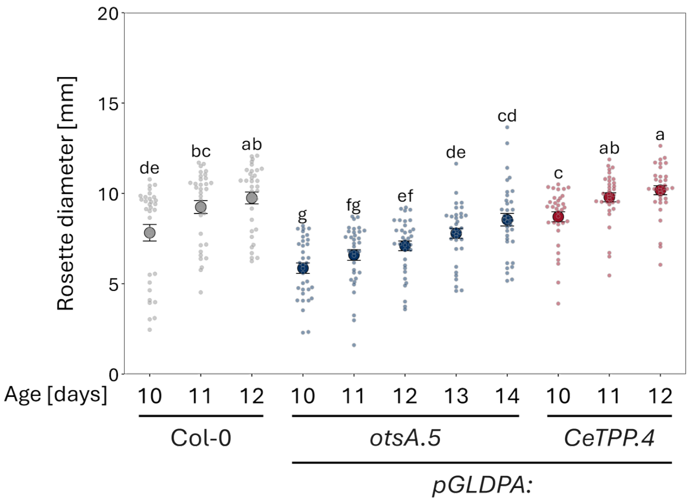


**Figure S8 Changing vascular Tre6P levels alters the rosette diameter.** Rosette diameter of Arabidopsis wildtype (Col-0, grey), *pGLDPA:otsA.5* (high Tre6P in the vasculature, blue), and *pGLDPA:CeTPP.4* (low Tre6P in the vasculature, red) seedlings ) seedlings grown vertically on ½ MS medium scored over time (age [days] = DAG). Displayed are the mean ± SEM (Large dots and error bars), with individual datapoints being shown as small dots ($n \geq 33$). Letters indicate significant differences between genotypes according to a one-way ANOVA with *post hoc* LSD testing ($p < 0.05$).

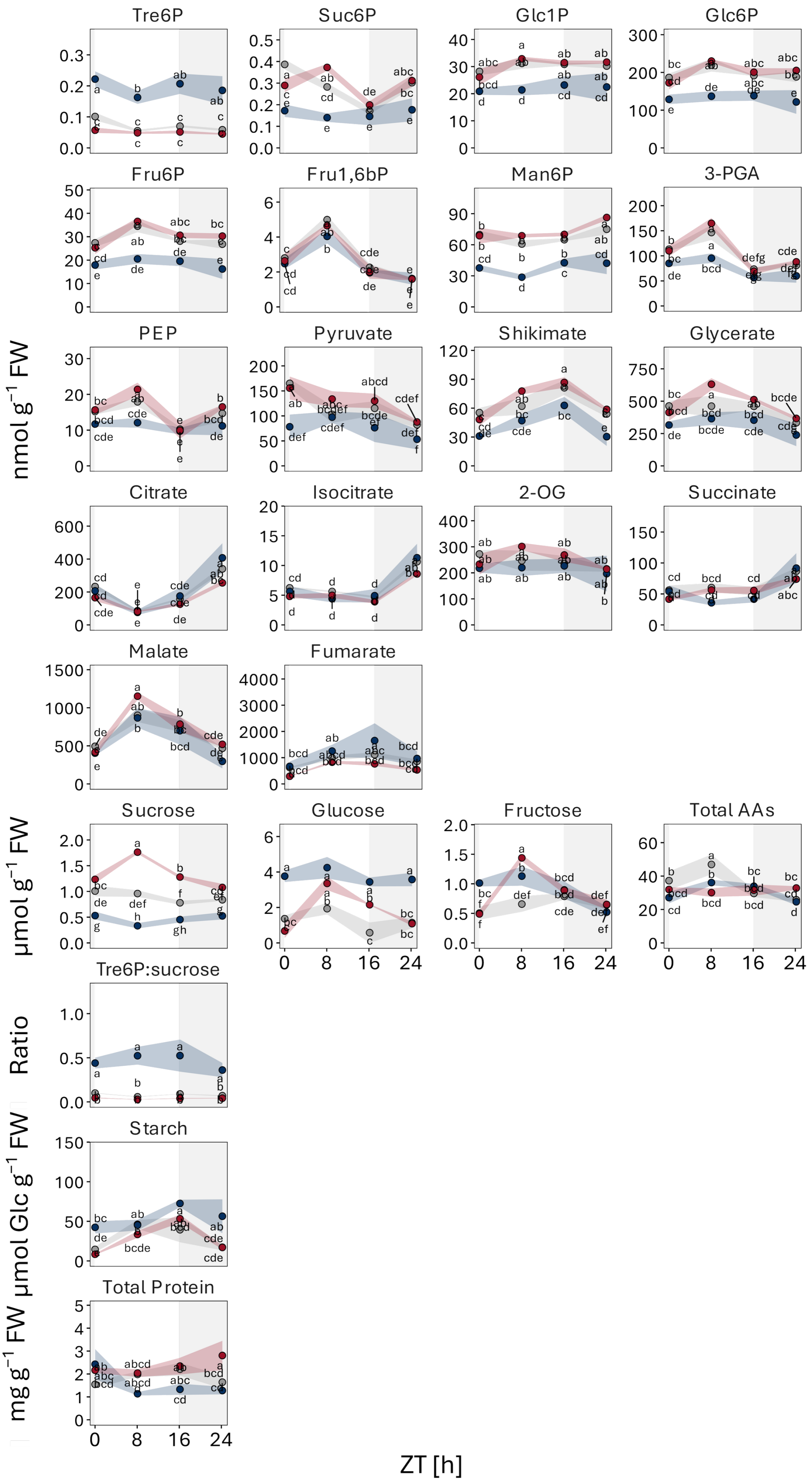
Tre6P
Suc6P
Glc1P
Glc6P
Fru6P
Fru1,6bP
Man6P
3-PGA
PEP
Pyruvate
Shikimate
Glycerate
Citrate
Isocitrate
2-OG
Succinate
Malate
Fumarate
Sucrose
Glucose
Fructose
Total AAs
Tre6P:sucrose
Starch
Total Protein
nmol g$^{-1}$ FW
μmol g$^{-1}$ FW
Ratio
μmol Glc g$^{-1}$ FW
mg g$^{-1}$ FW
ZT [h]

**Figure S9 Metabolite levels over the course of a day in Arabidopsis lines with altered Tre6P levels in the vasculature.** a) Analysis of Tre6P and other metabolite levels in the shoots of lines expressing a heterologous TPS (otsA) or TPP (CeTPP) under the control of a vasculature-specific promoter. Shoots and roots of wild-type (Col-0, grey), *pGLDPA:otsA.5* (more Tre6P in the vasculature, blue), *pGLDPA:CeTPP.4* (less Tre6P in the vasculature, red) seedlings were harvested right after dawn (zeitgeber time (ZT) 0), 8 h after dawn (ZT8), immediately after the onset of darkness (ZT16), and right before dawn (ZT24) from plants grown in a 16-h photoperiod (100 µmol $m^{-2}$ $s^{-1}$ irradiance) with 22°C day/18°C night temperatures 10 DAG. Metabolite levels are displayed as mean ± SEM ($n \geq 3$). Letters indicate significant differences between genotypes according to a one-way ANOVA with post hoc LSD testing ($p < 0.05$).

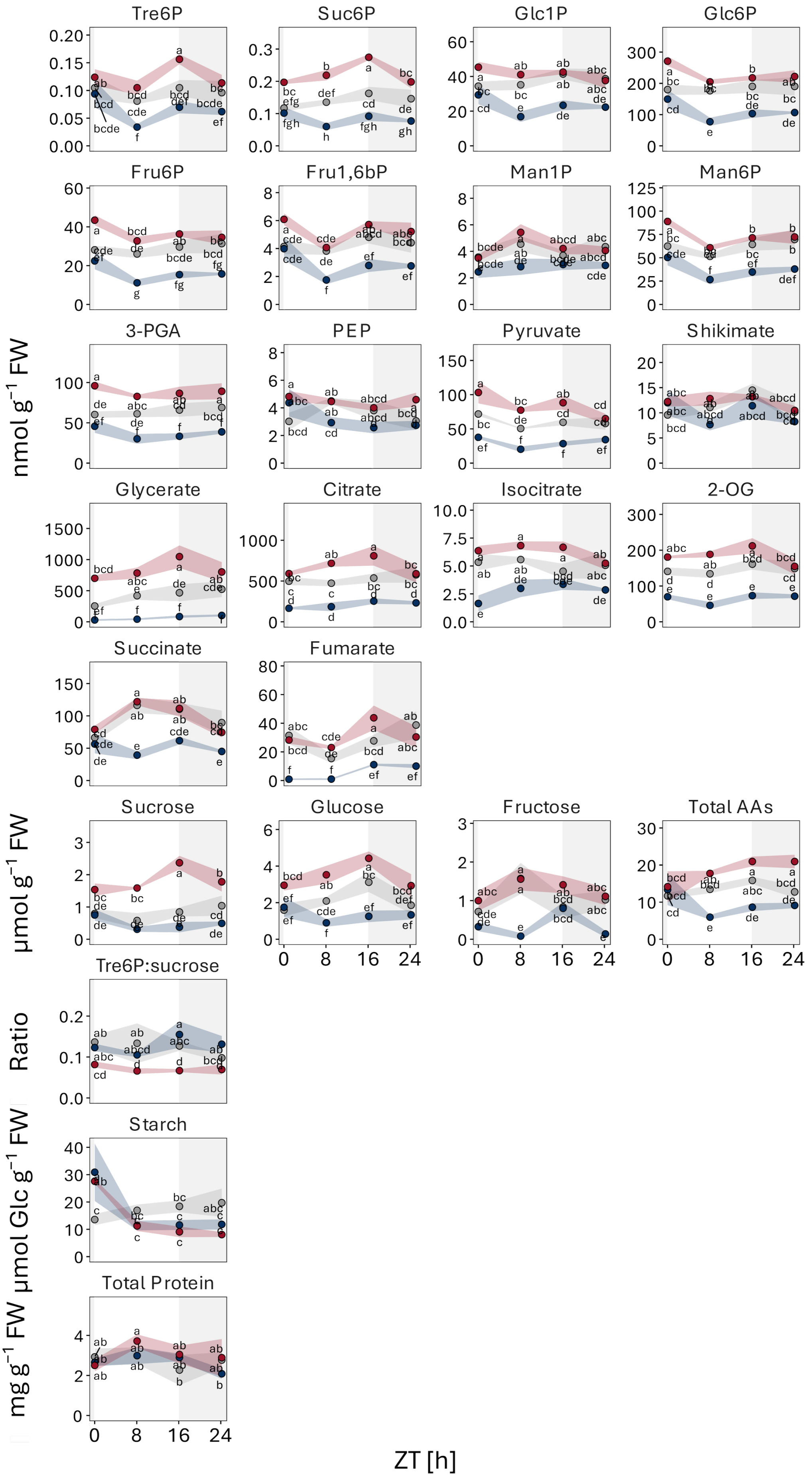
Tre6P
Suc6P
Glc1P
Glc6P
Fru6P
Fru1,6bP
Man1P
Man6P
3-PGA
PEP
Pyruvate
Shikimate
Glycerate
Citrate
Isocitrate
2-OG
Succinate
Fumarate
Sucrose
Glucose
Fructose
Total AAs
Tre6P:sucrose
Starch
Total Protein
nmol g⁻¹ FW
µmol g⁻¹ FW
Ratio
µmol Glc g⁻¹ FW
mg g⁻¹ FW
ZT [h]

**Figure S9 continued.** b) Analysis of Tre6P and other metabolite levels in the roots of lines expressing a heterologous TPS (otsA) or TPP (CeTPP) under the control of a vasculature-specific promoter. Shoots and roots of wild-type (Col-0, grey), *pGLDPA:otsA.5* (more Tre6P in the vasculature, blue), *pGLDPA:CeTPP.4* (less Tre6P in the vasculature, red) seedlings were harvested right after dawn (zeitgeber time (ZT) 0), 8 h after dawn (ZT8), immediately after the onset of darkness (ZT16), and right before dawn (ZT24) from plants grown in a 16-h photoperiod (100 µmol $m^{-2}$ $s^{-1}$ irradiance) with 22°C day/18°C night temperatures 10 DAG. Metabolite levels are displayed as mean ± SEM ($n \geq 3$). Letters indicate significant differences between genotypes according to a one-way ANOVA with post hoc LSD testing ($p < 0.05$).

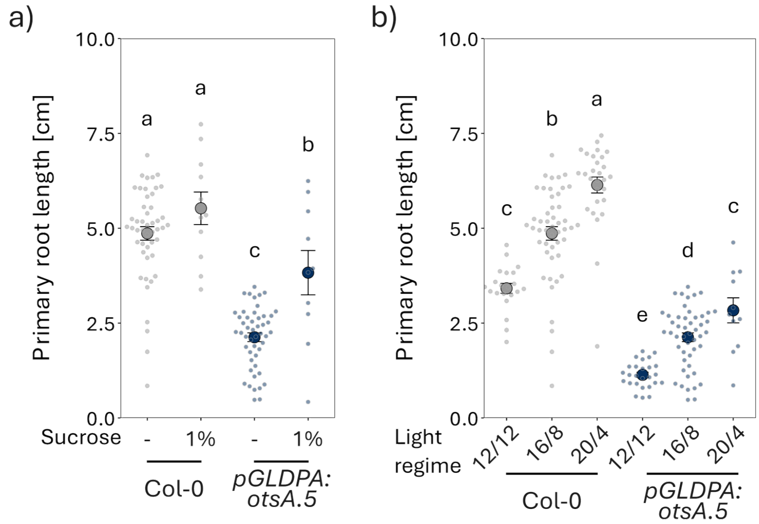


**Figure S10 Supplying carbon alone cannot rescue the root growth defects of lines with higher Tre6P in the vasculature.** Root morphology of Arabidopsis lines expressing a bacterial TPS (otsA) under the control of a vasculature-specific promoter supplied with excess carbon. a) Primary root length of wild-type (Col-0, grey) and *pGLDPA:otsA.5* (more Tre6P in the vasculature, blue) seedlings grown vertically on ½ MS medium or on ½ MS medium supplemented with 1% (w/v) sucrose in a 16-h photoperiod (100 µmol $m^{-2}$ $s^{-1}$ irradiance) with 22°C day/18°C night temperatures 10 DAG. b) The same lines grown on ½ MS medium in different photoperiods (ranging from 12 to 20 hours of light per day). Displayed are the mean ± SEM (Large dots and error bars), with individual datapoints being shown as small dots (a) n ≥ 10; b) n ≥ 11). Letters indicate significant differences between treatments according to a one-way ANOVA with *post hoc* LSD testing ($p < 0.05$).

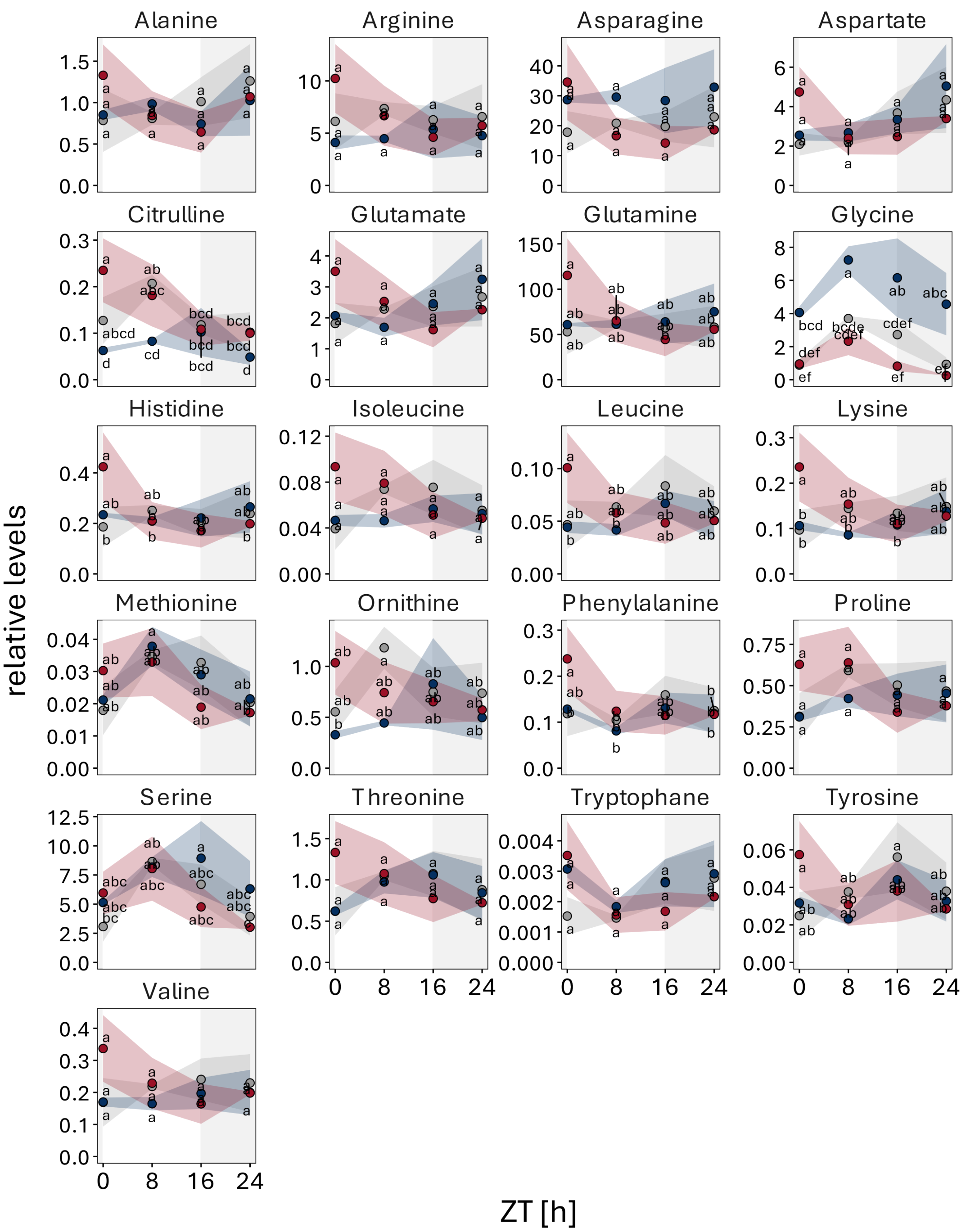


**Figure S11 Individual amino acid levels over the course of a day in Arabidopsis lines with altered Tre6P levels in the vasculature.** a) Analysis of individual amino acid levels in the shoots of lines expressing a heterologous TPS (otsA) or TPP (CeTPP) under the control of a vasculature-specific promoter. Shoots and roots of wild-type (Col-0, grey), *pGLDPA:otsA.5* (more Tre6P in the vasculature, blue), *pGLDPA:CeTPP.4* (less Tre6P in the vasculature, red) seedlings were harvested right after dawn (zeitgeber time (ZT) 0), 8 h after dawn (ZT8), immediately after the onset of darkness (ZT16), and right before dawn (ZT24) from plants grown in a 16-h photoperiod (100 µmol $m^{-2}$ $s^{-1}$ irradiance) with 22°C day/18°C night temperatures 10 DAG. Metabolite levels are displayed as mean ± SEM (n ≥ 3). Letters indicate significant differences between genotypes according to a one-way ANOVA with post hoc LSD testing ($p < 0.05$).

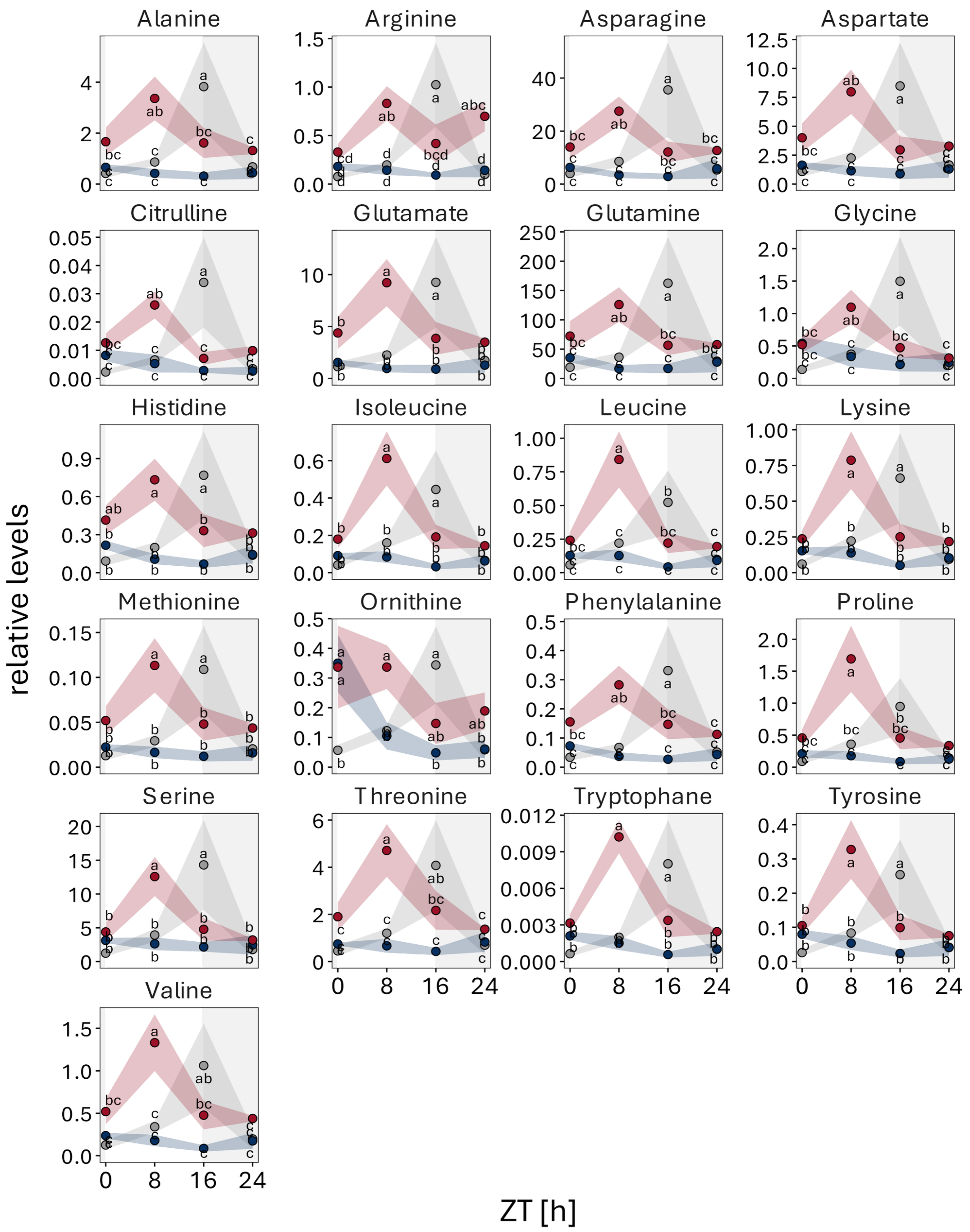


**Figure S11 continued.** b) Analysis of individual amino acid levels in the roots of lines expressing a heterologous TPS (otsA) or TPP (CeTPP) under the control of a vasculature-specific promoter. Shoots and roots of wild-type (Col-0, grey), *pGLDPA:otsA.5* (more Tre6P in the vasculature, blue), *pGLDPA:CeTPP.4* (less Tre6P in the vasculature, red) seedlings were harvested right after dawn (zeitgeber time (ZT) 0), 8 h after dawn (ZT8), immediately after the onset of darkness (ZT16), and right before dawn (ZT24) from plants grown in a 16-h photoperiod (100 µmol $m^{-2}$ $s^{-1}$ irradiance) with 22°C day/18°C night temperatures 10 DAG. Metabolite levels are displayed as mean ± SEM (n ≥ 3). Letters indicate significant differences between genotypes according to a one-way ANOVA with post hoc LSD testing ($p < 0.05$).

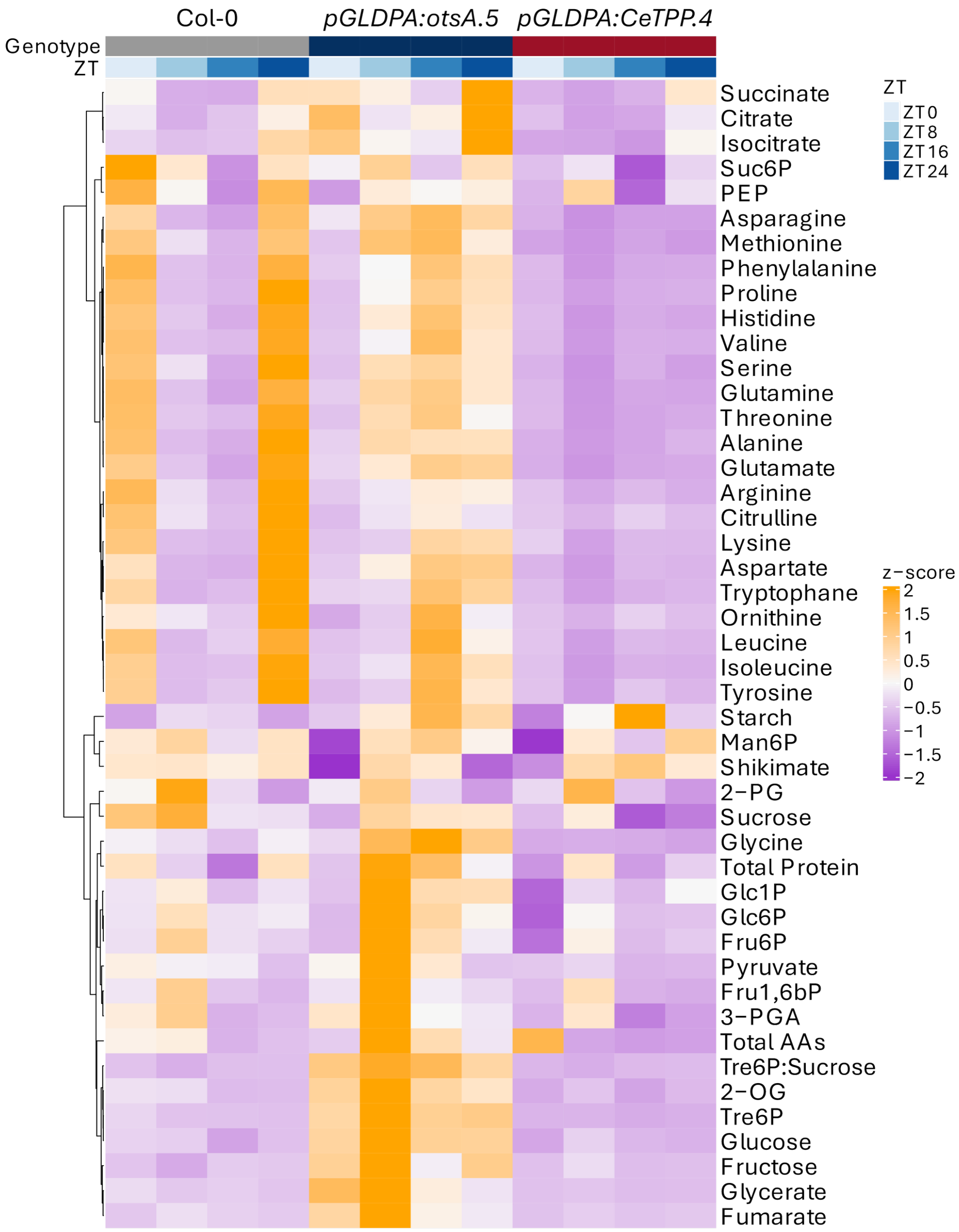


**Figure S12 Changing Tre6P levels alters metabolite allocation.** Hierarchical clustering analysis of Tre6P and other metabolite data from lines expressing a heterologous TPS (otsA) or TPP (CeTPP) under the control of a vasculature-specific promoter (data from S9, S11). Shoots (green) and roots (brown) of wild-type (Col-0, grey), *pGLDPA:otsA.5* (more Tre6P in the vasculature, blue), *pGLDPA:CeTPP.4* (less Tre6P in the vasculature, red) seedlings were harvested separately right after dawn (zeitgeber time (ZT) 0, very light blue), 8 h after dawn (ZT8, light blue), immediately after the onset of darkness (ZT16, mid blue), and right before dawn (ZT24, dark blue) from plants grown in a 16 h photoperiod (100 µmol $m^{-2}$ $s^{-1}$ irradiance) with 22°C day/18°C night temperatures 10 DAG. Shoot:root ratios of each metabolite were calculated and are presented as a heatmap, with high and low ratios represented in orange and purple, respectively. Hierarchical clustering of metabolites was performed using Pearson correlation to compute the distance matrix (1 – r), followed by Ward's minimum variance algorithm (*ward.D2*), and is represented by a dendrogram.

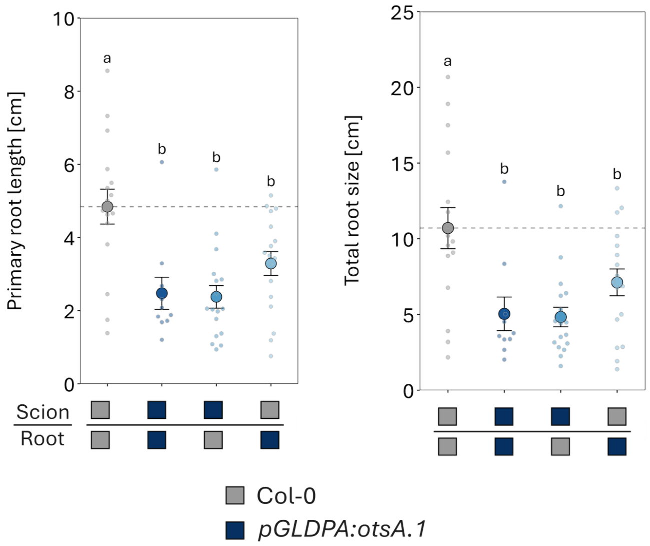


**Figure S13 Root morphology of grafted Arabidopsis seedlings with root or shoot vasculature specific expression of a bacterial TPS (otsA).** Primary (PR) and total root sizes (sum of the PR and all lateral root lengths) of grafted wildtype (Col-0, grey) and *pGLDPA:otsA.1* (high Tre6P in the vasculature, shades of blue represent different graft combinations) seedlings grown vertically on ½ MS medium. Displayed are the mean ± SEM (Large dots and error bars), with individual datapoints being shown as small dots (n ≥ 10). Letters indicate significant differences between genotypes according to a one-way ANOVA with post hoc LSD testing ($p < 0.05$).

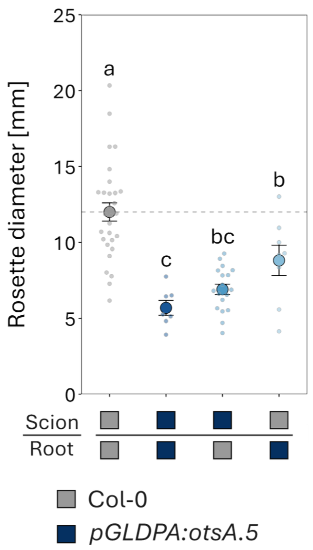


**Figure S14 The shoot growth in seedlings with alterations in Tre6P in the vasculature of shoots, or roots is consistent with their root growth phenotype.** Shoot morphology of grafted Arabidopsis seedlings with root or shoot vasculature specific expression of a bacterial TPS (otsA). Shoot diameter of grafted wildtype (Col-0, grey) and *pGLDPA:otsA.5* (high Tre6P in the vasculature, shades of blue represent different graft combinations) seedlings shown in Figure 6. Displayed are the mean ± SEM (Large dots and error bars), with individual datapoints being shown as small dots ($n \geq 7$). Letters indicate significant differences between genotypes according to a one-way ANOVA with *post hoc* LSD testing ($p < 0.05$).

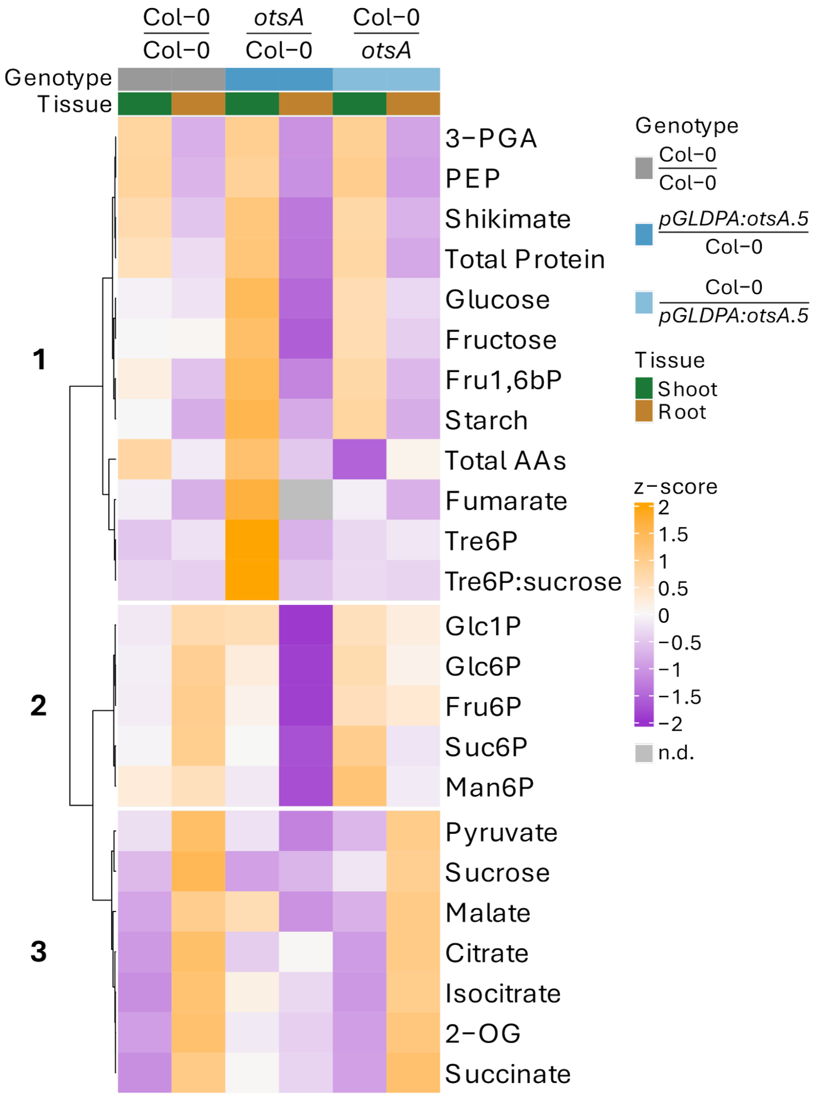


**Figure S15 Systemic changes in vascular Tre6P levels are accompanied by widespread changes in primary metabolism.** Hierarchical clustering analysis of Tre6P and other metabolite levels from grafted Arabidopsis seedlings with root or shoot vasculature-specific expression of a bacterial TPS (otsA) (data from Figure S16). Shoots and roots of grafted wildtype (Col-0) and *pGLDPA:otsA.5* (high Tre6P in the vasculature, shades of blue represent different graft combinations) seedlings 8 h after dawn from plants grown in a 16 h photoperiod (100 µmol m$^{-2}$ s$^{-1}$ irradiance) with 22°C day/18°C night temperatures 19 DAG. Z-scores of the mean were calculated for each metabolite and are presented as a heatmap, with high and low z-scores represented in orange and purple. Metabolites not detected in either tissue are shown in grey. Hierarchical clustering of metabolites was performed using Pearson correlation to compute the distance matrix (1 – r), followed by Ward's minimum variance algorithm (*ward.D2*), and is represented by a dendrogram with three distinct clusters.

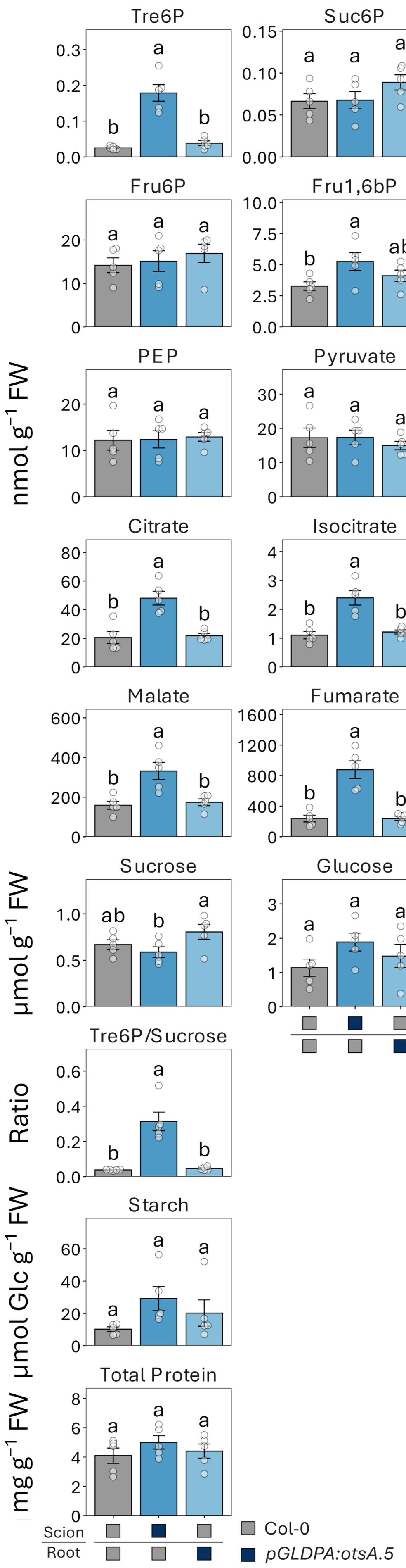
Tre6P
Suc6P
Glc1P
Glc6P
Fru6P
Fru1,6bP
Man6P
3−PGA
PEP
Pyruvate
Shikimate
Glycerate
Citrate
Isocitrate
2−OG
Succinate
Malate
Fumarate
Sucrose
Glucose
Fructose
Total AAs
Tre6P/Sucrose
Starch
Total Protein
nmol g$^{-1}$ FW
µmol g$^{-1}$ FW
Ratio
µmol Glc g$^{-1}$ FW
mg g$^{-1}$ FW
Scion
Root
Col-0
*pGLDPA:otsA.5*

**Figure S16 Tre6P systemically alters allocation of metabolites between shoots and roots.** a) Analysis of Tre6P and other metabolite levels in shoots of grafted Arabidopsis seedlings with root or shoot vasculature specific expression of a bacterial TPS (otsA). Shoots and roots of grafted wildtype (Col-0) and *pGLDPA:otsA.5* (high Tre6P in the vasculature, shades of blue represent different graft combinations) seedlings 8 h after dawn from plants grown in a 16 h photoperiod (100 µmol $m^{-2}$ $s^{-1}$ irradiance) with 22°C day/18°C night temperatures 19 DAG. Metabolite levels are displayed as mean ± SEM ($n \geq 4$). Letters indicate significant differences between genotypes according to a one-way ANOVA with post hoc LSD testing ($P \leq 0.05$). a) Shoot metabolite levels.

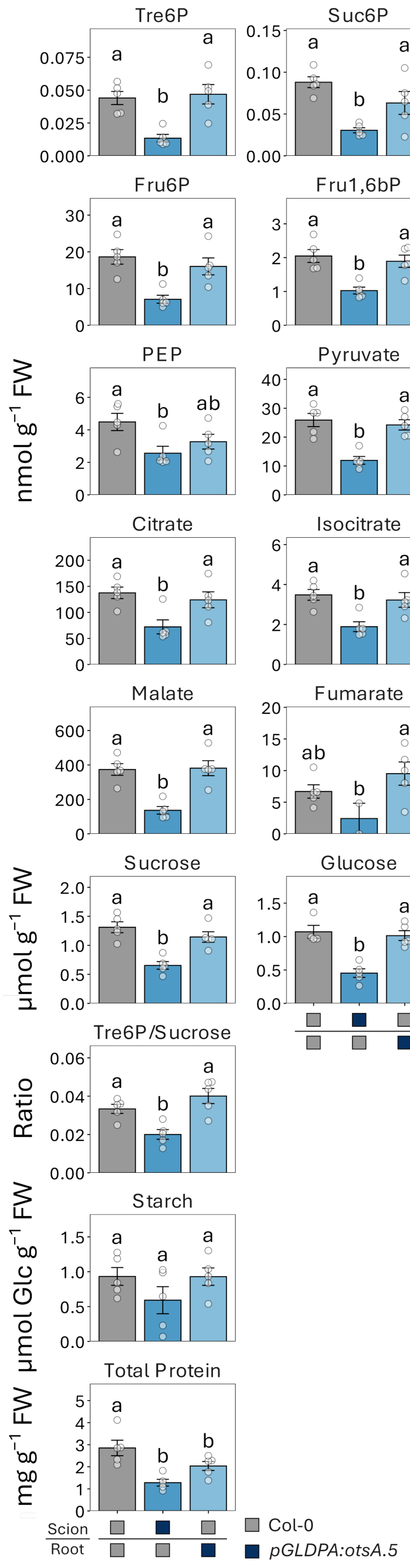

nmol g$^{-1}$ FW
Tre6P
Suc6P
Glc1P
Glc6P
Fru6P
Fru1,6bP
Man6P
3−PGA
PEP
Pyruvate
Shikimate
Glycerate
Citrate
Isocitrate
2−OG
Succinate
Malate
Fumarate
µmol g$^{-1}$ FW
Sucrose
Glucose
Fructose
Total AAs
Ratio
Tre6P/Sucrose
µmol Glc g$^{-1}$ FW
Starch
mg g$^{-1}$ FW
Total Protein
Scion
Root
Col-0
*pGLDPA:otsA.5*

**Figure S16 continued.** b) Analysis of Tre6P and other metabolite levels in roots of grafted Arabidopsis seedlings with root or shoot vasculature specific expression of a bacterial TPS (otsA). Shoots and roots of grafted wildtype (Col-0) and *pGLDPA:otsA.5* (high Tre6P in the vasculature, shades of blue represent different graft combinations) seedlings 8 h after dawn from plants grown in a 16 h photoperiod (100 µmol $m^{-2}$ $s^{-1}$ irradiance) with 22°C day/18°C night temperatures 19 DAG. Metabolite levels are displayed as mean ± SEM ($n \geq 4$). Letters indicate significant differences between genotypes according to a one-way ANOVA with post hoc LSD testing ($P \leq 0.05$). a) Shoot metabolite levels.

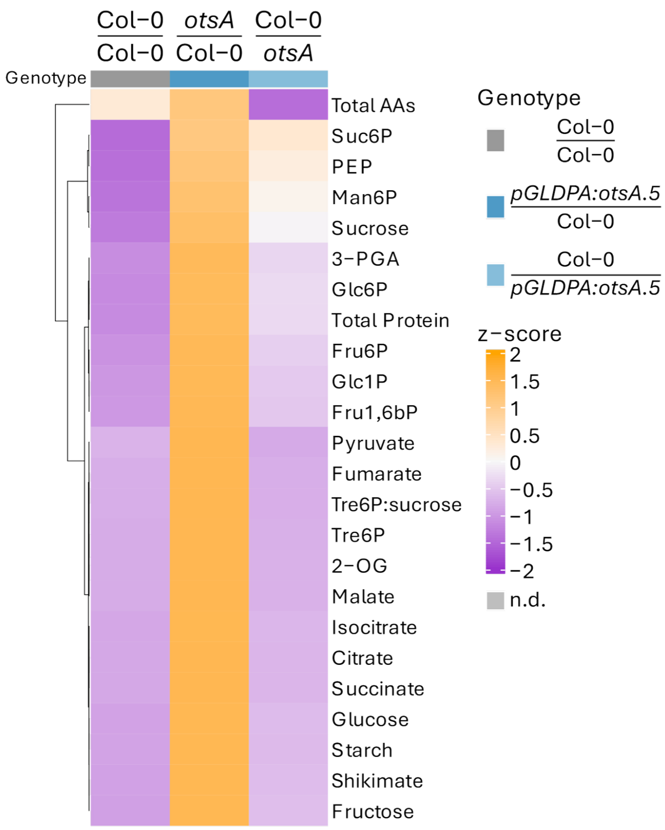


**Figure S17 Systemic changes in vascular Tre6P levels are accompanied by widespread changes in primary metabolism.** Hierarchical clustering analysis of Tre6P and other metabolite shoot to root ratios from grafted Arabidopsis seedlings with root or shoot vasculature-specific expression of a bacterial TPS (otsA). Shoots and roots of grafted wildtype (Col-0) and *pGLDPA:otsA.5* (high Tre6P in the vasculature, shades of blue represent different graft combinations) seedlings were harvested separately 8 h after dawn from plants grown in a 16 h photoperiod (100 µmol $m^{-2}$ $s^{-1}$ irradiance) with 22°C day/18°C night temperatures 19 DAG. Shoot:root ratios were calculated for each metabolite and are presented as a heatmap, with high and low ratios represented in orange and purple, respectively. Hierarchical clustering of metabolites was performed using Pearson correlation to compute the distance matrix (1 – r), followed by Ward's minimum variance algorithm (*ward.D2*), and is represented by a dendrogram.

**Table S1** single guide RNA (sgRNA) sequences used for the generation of the *CRISPR* $TPS1[\Delta N_{5\text{-}90}]$ line. The PAM sequences (not included in final constructs) are underlined.

| **sgRNA name** | **Sequence (5′ → 3′)** |
|---|---|
| TPS1_dN_sgRNA_1 | GCGCACGATGATGCGTGTGAG \| AGG |
| TPS1_dN_sgRNA_2 | GTTTTGGTGTGAGCGTATGCC \| TGG |

**Table S2** Oligonucleotide primers used for genotyping the deletions of the N-terminal domain in the *TPS1[ΔN]* lines.

| **Primer name** | **Sequence (5′ → 3′)** |
|---|---|
| TPS1_N-term_fwd | TTATTCCTTGGCCTGATGGGAC |
| TPS1_N-term_rvs | CCAAAGCTTTGCTAAGTGCCTT |
