## Supporting Method for "Systemic and local regulation of root growth by vascular trehalose 6-phosphate is correlated with re-allocation of primary metabolites between shoots and roots"

**Metabolite Analysis by IC–Orbitrap MS**

Metabolites were analyzed using a Dionex ICS-6000 HPIC system (Thermo Scientific) coupled to a Q Exactive Plus quadrupole-Orbitrap mass spectrometer (Thermo Scientific) according to Schwaiger et al., 2017 (DOI: 10.1021/acs.analchem.7b01624). The method is based on Lunn et al., 2006 (DOI: 10.1042/BJ20060083) and Figueroa et al., 2016 (DOI: 10.1111/tpj.13114). Samples (10 µL) were injected via a Dionex AS-AP autosampler in push-partial mode with a 25 µL loop; the autosampler was maintained at 4 °C.

Anion-exchange chromatography was on a Dionex IonPac AS11-HC column (2 mm × 250 mm, 4 µm; Thermo Scientific) with an AG11-HC guard column (2 mm × 50 mm, 4 µm) at 30 °C. A potassium hydroxide (KOH) gradient was generated by an eluent generator and delivered at 380 µL $min^{-1}$: 5 mM KOH for 1 min, ramped to 30 mM over 15 min, then to 95 mM over 5 min, held for 4 min, and reduced back to 5 mM for 10 min of re-equilibration. A Dionex AERS 600 suppressor (2 mm) in legacy mode at 20 °C exchanged $K^+$ for $H^+$, converting KOH to water and reducing salt load before ionization.

Because stable electrospray could not be achieved with water alone, a make-up flow of methanol with 10 mM acetic acid was introduced using an AXP pump at 150 µL $min^{-1}$ via a tee-junction immediately before the ESI source. Negative-mode ionization was applied with the following parameters: sheath gas 30, auxiliary gas 20, sweep gas 0, spray voltage –2.8 kV, capillary temperature 230 °C, S-Lens RF 45, and auxiliary gas heater 380 °C.

Data were acquired in full-scan mode over an m/z range of 70–900 at a resolution of 140,000 (at m/z 200), with an automatic gain control (AGC) target of $3 \times 10^6$ and a maximum injection time (IT) of 200 ms.

Between 9–15 min of the chromatographic run, a single-ion monitoring (SIM) experiment targeting trehalose-6-phosphate (m/z 421.07527) and its $^{13}C$-labeled isotopologue (m/z 433.07527) was performed in parallel, at a resolution of 70,000 with an AGC target of $1 \times 10^6$ and a maximum IT of 200 ms.

**Generation of an internal standard for Tre6P**

A uniformly $^{13}C$-labeled trehalose-6-phosphate internal standard was synthesized as described by Lunn et al. (2006) using $^{13}C$-labeled UDP-glucose and $^{13}C$-labeled glucose 6-phosphate. In brief, 10 mM $^{13}C$-Glc6P and 10 mM $^{13}C$-UDP-Glc were incubated for 3 h at 30°C in a buffer consisting of 25 mM HEPES, 5 mM MgCl2, 0.5 mM EDTA (pH 7.2), and 5 U of purified OtsA protein. OtsA proteins were denatured by incubating the reaction mix for 2 min at 99°C.

Samples were centrifuged at 2,200 g at 15°C. Standards were filtered through nitrocellulose membrane filters.

**Data Analysis and Calibration**

Data were processed and quantified using Skyline (DOI: 10.1021/acs.jproteome.9b00640). Calibration curves were generated using authentic reference standards and were normalized with the corresponding $^{13}C$-labeled internal standards (malate, fumarate, citrate, phosphoenolpyruvate, trehalose 6-phosphate, glucose 6-phosphate, glucose 1-phosphate and UDP-glucose). Each calibration curve consisted of six points, prepared by serially halving the highest concentration. To ensure accuracy and reproducibility, three intermediate quality-control (QC) samples were included within the calibration range.
